# Oncolytic Reovirus mediates innate-driven SARS-CoV-2 elimination in the absence of cell toxicity

**DOI:** 10.64898/2025.12.17.694833

**Authors:** Samantha Garcia-Cardenas, Gemma Swinscoe, Amy Moran, Liam Barningham, Graham Cook, Adel Samson, Sam Wilson, Michael Malim, Russell Hughes, Stephen Griffin

## Abstract

Interplay between type I interferon (IFN) driven innate responses and viral antagonism strongly influences SARS-CoV-2 transmission and the COVID-19 disease course. Hence, variant adaptation includes diminished induction of IFN stimulated genes (ISG) and/or evasion of their effector functions. Exogenous IFN treatment “rewires” innate responses to drive virus elimination, yet therapeutic trials to date have been unremarkable. Resolving this paradox could translate to variant-agnostic innate immunotherapy.

By contrast, oncolytic viruses (OV) exhibit profoundly attenuated innate antagonism, resulting in potent IFN responses despite the inherently immunosuppressive nature of tumour microenvironments. Moreover, OV only undergo lytic replication within innate-deficient malignant cells, and not in cells where sufficient innate responses exist. This, combined with previous studies showing that OV suppressed replication of underlying oncogenic viruses in tumours, we explored whether clinical grade oncolytic *Orthoreovirus* (Reo) superinfection could eliminate SARS-CoV-2 from immune-competent lung epithelial cell lines in the absence of toxicity. Reo exerted profound activation of innate responses, including when SARS-CoV-2 infection was already established, rewiring cells towards an antiviral state emulating that of Reo infection alone. Both intracellular and paracrine mechanisms induced ISG repertoires including multiple known anti-SARS-CoV-2 effectors, as well as others that remain unvalidated. Amongst these, we demonstrate the first direct evidence that MX2 and XAF1 restrict SARS-CoV-2 replication. Thus, with an excellent safety record, self-amplification, and respiratory tract tropism, we propose that Reo superinfection may provide a tractable alternative to recombinant cytokines for innate antiviral immunotherapy.

## Introduction

The Coronavirus disease 2019 (COVID-19) pandemic, caused by zoonotic severe acute respiratory syndrome Coronavirus type 2 (SARS-CoV-2) [1, 2], has inflicted profound acute and long-term physical and socioeconomic harm. Along with SARS-CoV-1 [3], Middle East respiratory syndrome coronavirus (MERS-CoV) [4], and circulating bat CoVs with pandemic potential, the threat posed to human health by these viruses makes understanding their biology and development of new pan-CoV therapeutics a priority.

SARS-CoV-2 transmission, replication, and disease severity are strongly influenced by innate immunity, notably type I/III interferons (IFN) and multiple effector IFN stimulated genes (ISG) [5]. Allelic polymorphisms related to ISG repertoires or functions can dictate disease outcomes [6–8], as can the development of anti-IFN autoantibodies [9–11]. Hence, emergent viral variants evolve improved innate immune antagonism as they better-adapt to humans [12–14]. However, to date, recombinant type I/III IFN trials have failed to meet clinical trial primary endpoints for acute COVID-19 [15–17]. Resolving this paradox could lead to improved translational opportunities.

“Oncolytic viruses” (OV) [18] exhibit profound natural, or engineered attenuation of their ability to antagonise innate immune responses; this, in-part, determines selective lytic replication in malignant cells. However, OV also stimulate responses from normal tissues and tumour stroma, but their replication is profoundly limited as a result. OV therefore comprise potent innate adjuvants, able to switch tumours from immunologically “cold” to “hot” in the absence of toxicity in normal tissue. Whilst many OV are genetically modified, human *Orthoreovirus* Type 3, Dearing strain, (specifically the T3D^PL^, “Patrick Lee” Reovirus strain, designated herein: “Reo”) naturally exhibits such characteristics [19], and clinical grade virus (“Pelareorep”, Oncolytics Biotech, Calgary, CA) has fast-track FDA approval for advanced breast and pancreatic cancer.

Previously, we discovered that Reo exerted combined antiviral and anti-tumour effects in models of virus-driven cancer [20]. Hence, we asked whether it could rewire SARS-CoV-2 innate responses in SARS-CoV-2 models that are immunocompetent, yet do not mount a robust-enough response to clear the virus in the absence of exogenous IFN. Reo superinfection of “A549-AT” respiratory epithelial cells prompted primarily IFN-driven elimination of SARS-CoV-2 in the absence of toxicity. Transcriptomics confirmed progressive dominance of Reo, leading to rewiring of cellular gene expression over time, including when SARS-CoV-2 infection was already present. Reo-induced ISGs included multiple factors previously identified as restricting SARS-CoV-2 replication. However, focusing upon the early therapeutic response and correlating protein expression with transcript, we established the first direct experimental evidence that MX2 and XAF1 can restrict SARS-CoV-2 replication.

We therefore propose that further preclinical and translational exploration of Reo as an alternative format for innate antiviral immunotherapy is warranted. As its name suggests, Reo (“respiratory-<u>e</u>nteric <u>o</u>rphan”) exhibits a highly favourable safety profile which, combined with self-amplification in tissues and inherent respiratory and gastrointestinal tropism, lends itself towards a safe alternative platform for future repurposing for SARS-CoV-2, or other pathogenic viruses.

## Results

### Reo infects and promotes robust innate responses in lung epithelial cell lines

Workhorse respiratory epithelial cell lines such as A549-AT and Calu-3 have proven invaluable tools for characterising SARS-CoV-2 replication. However, A549 cells and derivatives fail to mount effective IFN responses to SARS-CoV-2, yet they respond to exogenous cytokine to mediate virus elimination. By contrast, productive Reo infection of both A549 (Figure 1, Figure S1, S2) and Calu-3 (Figure S3a-c) led to robust detection of Reo PAMP. However, replication persisted in spite of innate responses, and infectious virions were released from A549 cells, Figure (1a). Reo was unable to form plaques in A549, consistent with low cytotoxicity, and this was also evident from confluency measurements (Figure S1). Nevertheless, Reo and *uv*-inactivated Reo (*uv*-Reo) elicited robust innate responses from both cell types, with the latter response being more rapid but less durable, resulting in reduced cytokine production (Figure 1c, d). This difference may be attributable to the complete lack of viral innate antagonist proteins compared with the live T3D virus, which, albeit attenuated, do delay and dampen IFN responses to a degree [21].

**Figure 1.**
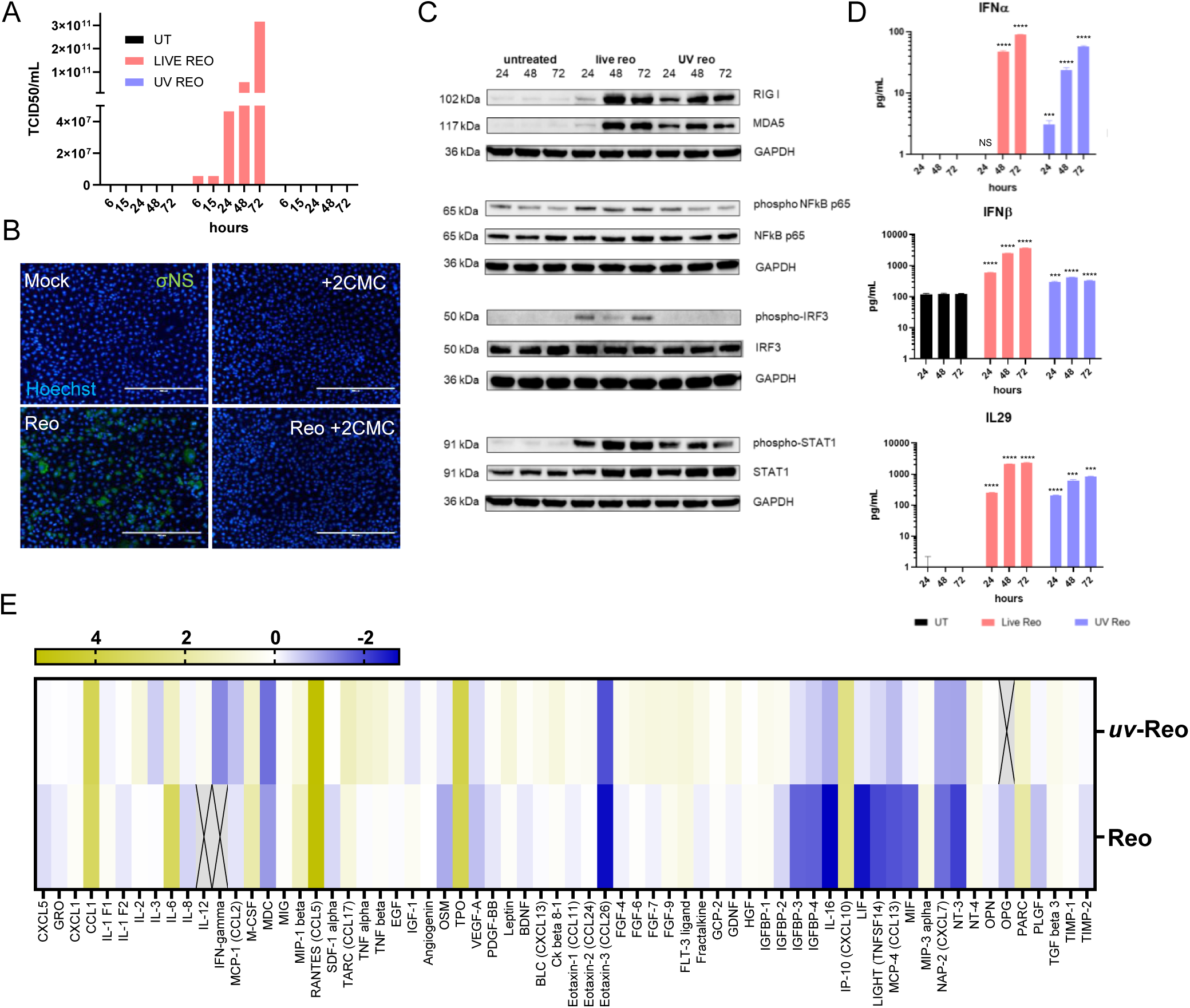
Response to Reo and uv-Reo by A549-AT cells. **A.** Output titres over time (24, 48, 72 hr) from infectious supernatants generated following infection of A549-AT cells with Reo virus at an M.O.I. of 0.1 Pfu/cell. Titres determined by TCID_50_ assay on L929 cells (n=1). **B.** Specific inhibition of Reo replication using 2CMC (30 μM) with visualisation of productively infected cells by immunofluorescent detection of σNS protein. **C.** Sensing of Reo or *uv*-Reo and pathway activation over time (24, 48, 72 hr post-stimulation) analysed by western blot. Top panel: induction of RIG-I and MDA-5 expression; second panel: induction of phosphorylated NFkB (p65); third panel: phosphorylated IRF3 induction; bottom panels: STAT1 phosphorylation. Gels are representative of two or three independent biological repeats (except p-IRF3 where n=1). **D.** Filtered supernatant corresponding to stimulation conditions in **C** were tested for type I/III IFN concentration by specific ELISA. Data represent the mean ± SD of three biological replicates and were analysed using ordinary two-way ANOVA with Tukey’s multiple comparisons test (*P < 0.05, **P < 0.01, ***P < 0.001, ****P < 0.0001). **E.** Determination of additional supernatant cytokine/chemokines and other factors by protein absorption shown as a heatmap plotting protein expression as -Log2 fold-change compared to controls. Cross-hatched blocks represent protein levels beneath the threshold of detection.

Reo infection of A549 cells elicited significant activation of IRF3, NFκB (p65), and STAT1 by 24 h post infection (hpi), indicative of pattern recognition and IFN production. Moreover, ISG induction was evident by markedly increased expression of RIG-I (*DDX58*) and MDA-5 (*IFIH1*) ISG by 48 hpi (Figure 1c). By contrast, *uv*-Reo activated IRF-3 in Calu-3, but not in A549-AT at the timepoints tested (Figure 1c, Figure S3a). Reo induced secretion of IFNα, IFNβ and IL29 from A549 cells, yet Calu-3 cells did not produce measurable IFNα, consistent with low mRNA expression in previous studies (Figure 1d, Figure S3b) [22]. Moreover, Reo/*uv*-Reo conditioned media ((*uv*)RCM) contained additional cytokines and chemokines, including CCL1, CCL5, CCL17, IL6, and CXCL10 (Figure 1e, Figure S3e, f). *uv*-RCM did not induce appreciable levels of IL6, whilst CCL17 was doubled (Figure 1e). Interestingly, recombinant CCL1 exerted anti-SARS-CoV-2 activity, albeit at high doses (Figure S3f).

### (uv)-Reo superinfection eliminates SARS-CoV-2 from cell culture

We next investigated Reo superinfection of A549-AT cells prior (−12 hr), during (0 hr) or after (+6 hr) infection with SARS-CoV-2, assessing the number of SARS-CoV-2 positive cells at 24-72 hr post infection (hpi) (Figure 2a, b). At 24 hpi, (*uv*)-Reo pre-treatment and higher multiplicity infection had a greater impact upon SARS-CoV-2 infection (Figure 2b, c), but at later times a profound reduction of SARS-CoV-2 infected cells was measurable under all conditions, with elimination of SARS-CoV-2 by 48 hpi when cells were pretreated (Figure 2d). We also obtained orthogonal confirmation that Reo replication was dispensable for short-term inhibition of SARS-CoV-2 infection using a nucleotide-based RNA dependent RNA polymerase (RdRP) inhibitor, 2-C-methylcytidine (2CMC), that we discovered to specifically block Reo replication but not SARS-CoV-2 (Figure 2e, Figure S3d, S5). Finally, immunofluorescence staining of superinfection experimental cultures revealed that Reo/SARS-CoV-2 dual-infected cells were a relatively rare occurrence, and where they did occur, Reo expression was generally lower, despite a much higher MOI (Figure 3). Given that SARS-CoV-2 was eventually eliminated by Reo/*uv*-Reo therapy, we surmise that coinfected cells will eventually become overtaken by Reo.

**Figure 2.**
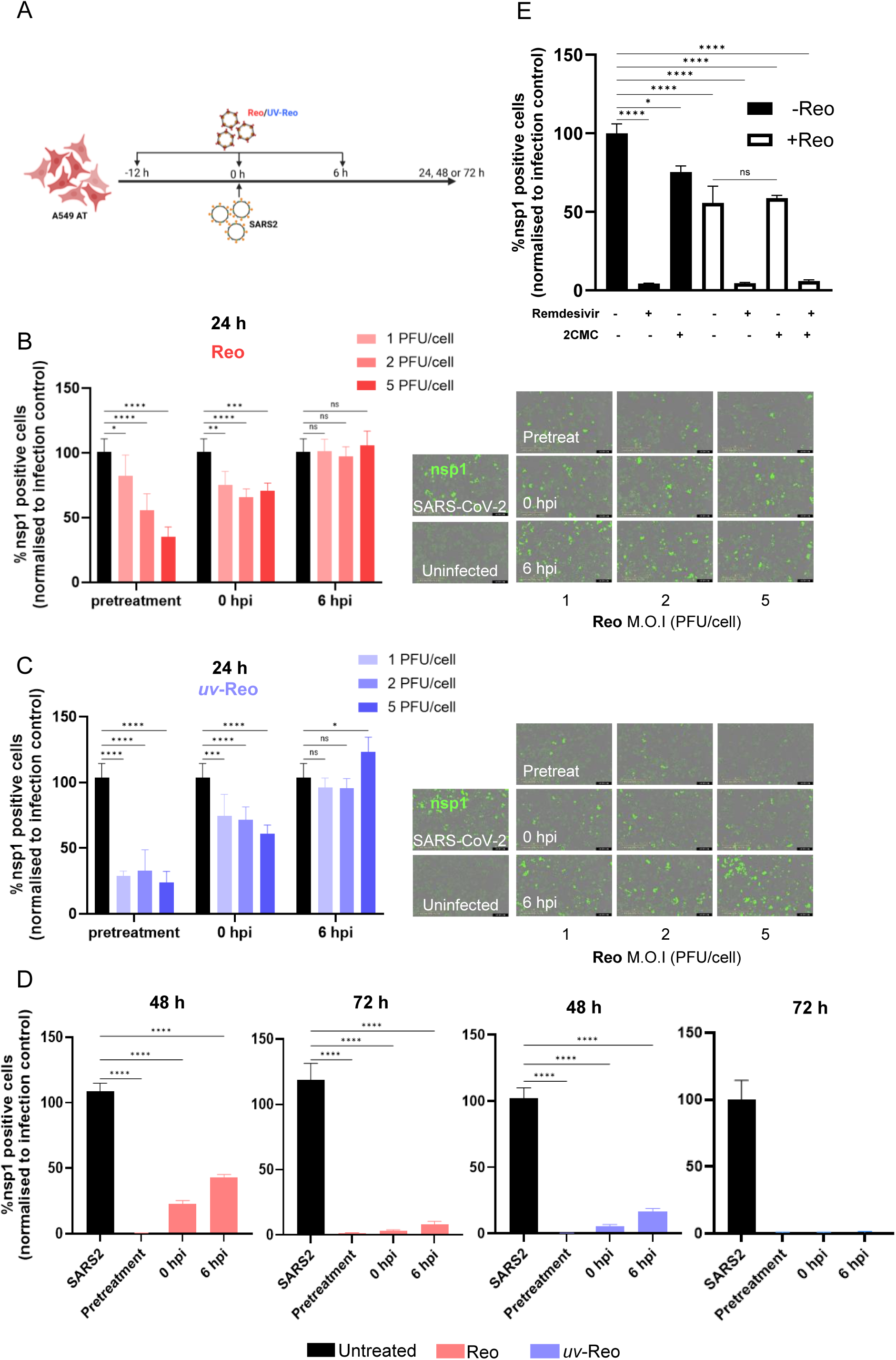
Short- and long-term responses to Reo and uv-Reo superinfection in A549-AT cells harbouring SARS-CoV-2. **A.** Schematic of experimental setup. **B.** NSP1 positive cells 24 hpi with SARS-CoV-2 (Eng/2) following mock (black bars) or Reo treatment at increasing M.O.I. (shades of red) either prior, coincident or following the 0-hr time point, as depicted in A. Right hand panels show representative IncuCyte images. Data represent the mean ± SD of three biological replicates and were analysed using ordinary two-way ANOVA with Tukey’s multiple comparisons test (*P < 0.05, **P < 0.01, ***P < 0.001, ****P < 0.0001). **C**. Data represented as per B, only following treatment with *uv*-Reo (shades of blue) in place of live virus. **D**. NSP1 positive cells were quantified at 48 and 72 hpi with Reo/*uv*-Reo treatment. Data represent the mean ± SD of three biological replicates and were analysed by using ordinary one-way ANOVA with Tukey’s multiple comparisons test (*P < 0.05, **P < 0.01, ***P < 0.001, ****P < 0.0001). **E**. NSP1 positive cells at 24 hpi with (white bars) or without (black bars) Reo treatment (0-hr), combined with selective inhibition of either SARS-CoV-2 (Remdesivir, 5 μM) or Reo (2CMC, 10 μM) replication. Data represent the mean ± SD of three experimental repeats and were analysed using ordinary multiple t-tests (*P < 0.05, **P < 0.01, ***P < 0.001, ****P < 0.0001).

**Figure 3.**
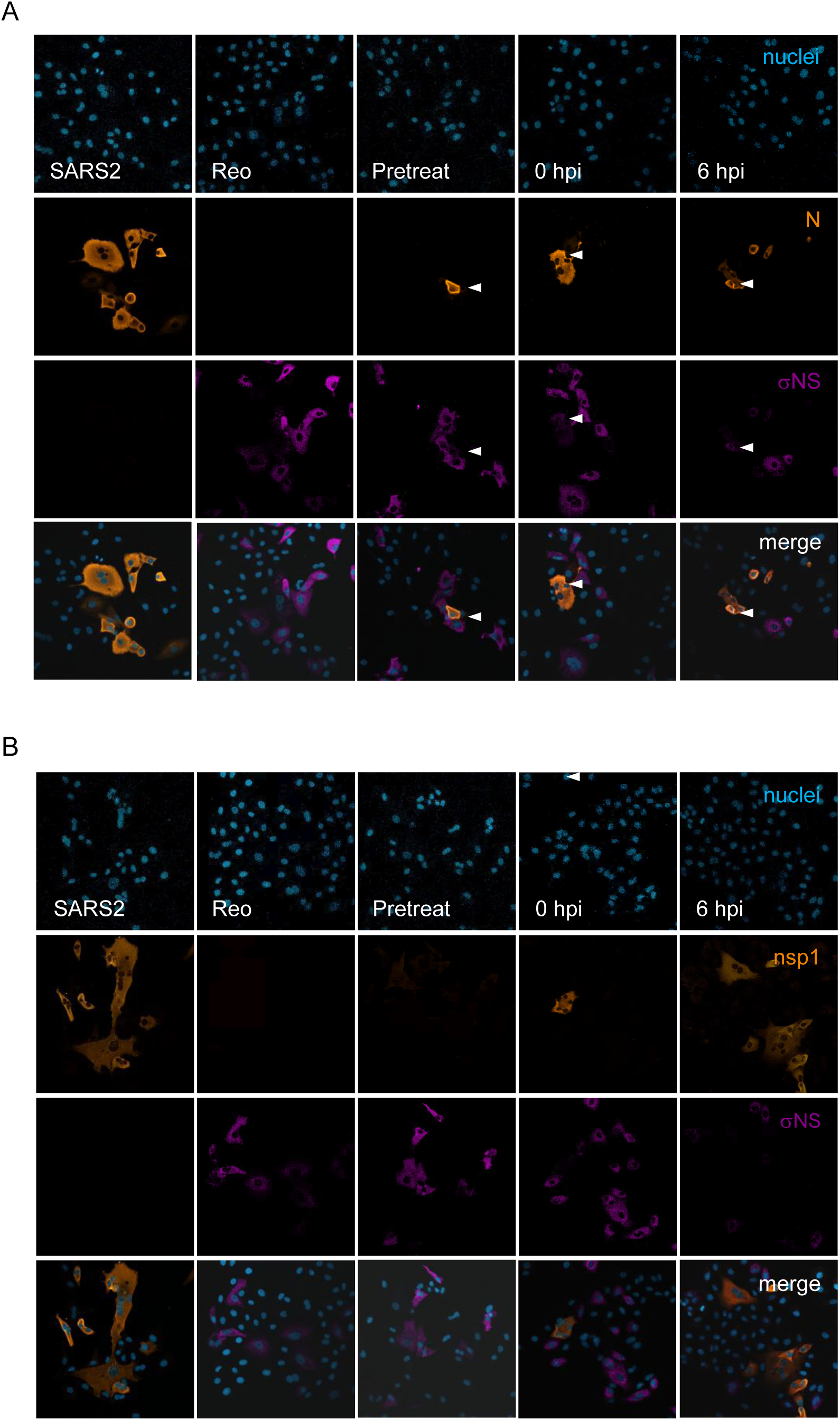
Reo and SARS-CoV-2 infected A549-AT cells during superinfection. Single and co-infected cultures grown in 96-well plates were fixed at 24 hpi then immunostained for SARS-CoV-2 and Reo antigens. **A.** Cells co-stained for SARS-CoV-2 N protein, Reo σNS, and counterstained with Hoechst. White arrows indicate co-stained cells. **B.** As per A, but SARS-CoV-2 NSP1.

### Reo superinfection exerts IFN-driven bystander antiviral effects

(*uv*-)Reo-conditioned media (RCM), filtered to remove Reo particles, was used to discriminate paracrine/bystander versus intracellular effects upon anti-SARS-CoV-2 potency (Figure 4a). Vero-T cells were infected with SARS-CoV-2 to avoid endogenous IFN responses, then treated using diluted (5, 10, or 20 % v/v) RCM derived from A549 cells (Figure 4a). Both *uv*-RCM and RCM effectively suppressed SARS-CoV-2 replication at 24 hpi, although lower concentrations of *uv*-RCM were less effective at treating an established infectious culture consistent with lower cytokine content (Figure 4a, b, c). We then compared RCM efficacy to treatment with the main cytokine components, namely IFNβ and IL-29, either individually or in combination. Notwithstanding differences in commercial biological activities, RCM was consistently more potent compared with individual or combined recombinant cytokines, with IFNβ more effective than IL-29 at the timepoint measured (Figure 4d). Finally, we compared RCM efficacy against the ancestral virus (Eng/2) versus the delta (B.1.617.2) and omicron (BA.2) SARS-CoV-2 variants of concern (VoC, Figure 4e, Figure S2b) at 24 hpi. Similar effects were observed for delta versus the Wuhan virus, although *uv*-RCM appeared less effective. Interestingly, Omicron BA.2 exhibited a considerable degree of resistance compared with the other variants, especially to the lower potency *uv*-RCM (Figure 4e). Finally, RCM responses were highly sensitive to Ruxolitinib, consistent with type I/III IFN driven stimulation of JAK1/2 dependent signalling (Figure 4f). Interestingly, control conditioned media (CM) from A549-AT cells enhanced the infection of Vero-T cells by SARS-CoV-2, and this was also sensitive to Ruxolitinib treatment. We presume that CM contains factors in addition to normal serum that activate JAK-linked growth factor receptors, or other proliferative pathways.

**Figure 4.**
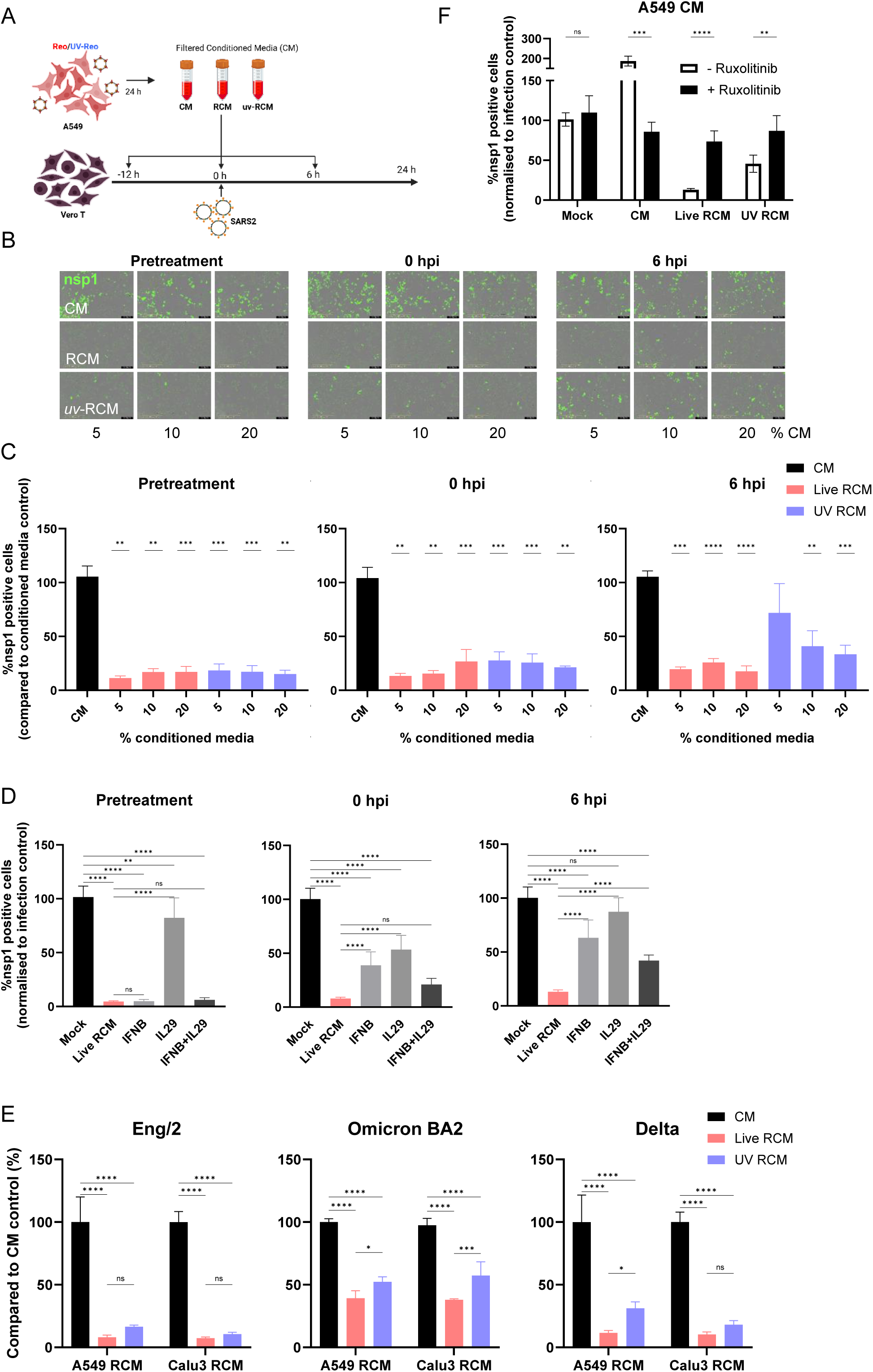
Reo and uv-Reo induced antiviral bystander effects are primarily, yet not completely IFN driven. **A.** Schematic of experimental plan comprising treatment of SARS-CoV-2 infected Vero-T cells using filtered A549 Reo, *uv*-Reo, or controlled conditioned media (RCM, *uv*-RCM, CM). **B.** Representative IncuCyte images of treated Vero-T cells 24 hpi following treatment prior, during or following infection, and at increasing proportions of RCM, *uv*-RCM, or CM. **C.** Quantification of NSP1 positive cells following treatments as described in B using CM (black), RCM (red), or *uv*-RCM (blue). Data represent the mean ± SD of three biological replicates and were analysed using ordinary multiple t-tests (*P < 0.05, **P < 0.01, ***P < 0.001, ****P < 0.0001). **D.** NSP1 positive cells were quantified at 24 hpi following RCM treatment (red bars), and compared to treatment with equivalent amounts of recombinant IFNβ, IL-29, or both (darkening shades of grey). Data represent the mean ± SD of three biological replicates and were analysed by using ordinary one-way ANOVA with Tukey’s multiple comparisons test (*P < 0.05, **P < 0.01, ***P < 0.001, ****P < 0.0001). **E.** Repeat of experiments in C comparing coincident treatment with RCM, *uv*-RCM, or CM (A549 or Calu-3 derived) versus Eng/2, Omicron BA.2, or Delta infection of Vero-T cells by NSP1 immunofluorescence. Data represent the mean ± SD of three biological replicates and were analysed using ordinary two-way ANOVA with Tukey’s multiple comparisons test (*P < 0.05, **P < 0.01, ***P < 0.001, ****P < 0.0001). **F**. Sensitivity of RCM, *uv*-RCM, or CM treatment of SARS-CoV-2 (Eng/2) infected cells at 24 hpi in the presence (black bars) or absence (white bars) of Ruxolitinib. Data represent the mean ± SD of three experimental repeats and were analysed using ordinary multiple t-tests (*P < 0.05, **P < 0.01, ***P < 0.001, ****P < 0.0001).

### Gene expression in SARS-CoV-2 infected cells is rewired following therapeutic superinfection

We used RNA sequencing (RNA-Seq) to analyse A549-AT cultures and interrogate the nature of the IFN/JAK dependent changes associated with Reo superinfection prior to, coincident with, or following SARS-CoV-2 infection. Samples taken at 24 hpi exhibited good separation between tightly clustered conditions on a three-dimensional PCA plot (Figure 5a).

**Figure 5.**
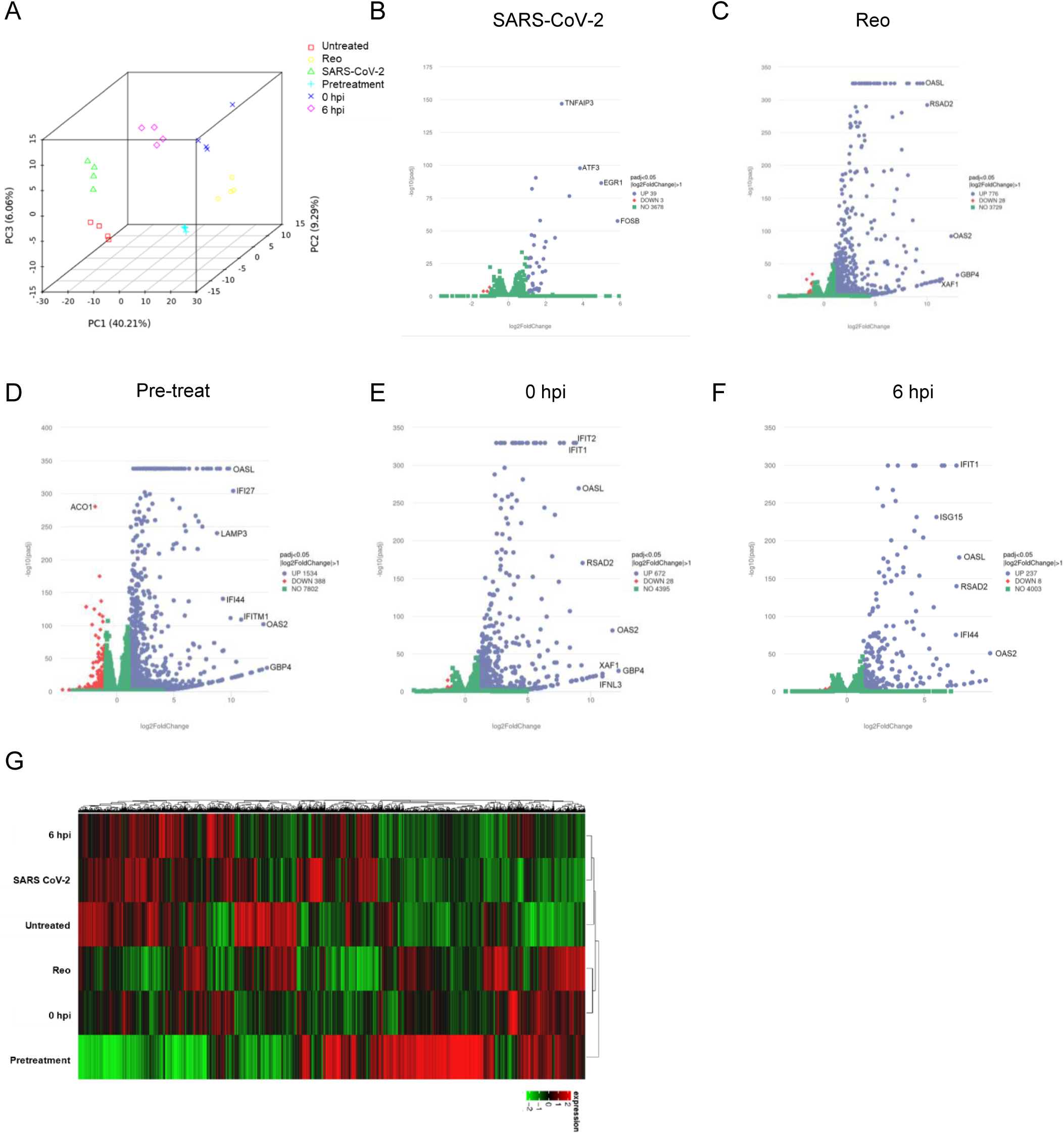
mRNA Sequencing (RNA-Seq) of SARS-CoV-2 infected A549-AT cultures following Reo superinfection. **A.** Three-dimensional principal component analysis of control, single infections and superinfection cultures. **B-F**. Volcano plots of DEGs from individual or superinfections compared with controls, plotting log_2_-fold change in gene expression versus - log_10_(P_adj_) values. Blue: significant, up-regulated; red: significant, down-regulated; green: non-significant, less than 2-fold change in expression. **G.** Heatmap of differential gene expression following hierarchical clustering analysis.

SARS-CoV-2 infection resulted in surprisingly low numbers of differentially expressed genes (DEG), with just 42 changes, of which all but two were upregulated (Figure 5b). Ontologically, these corresponded primarily to cell cycle and survival, with essentially no sign of innate immune interactions (Figure S9). Hence, the virus was able to infect cells and evade detection and/or antagonise the activation of PRRs and ensuing antiviral responses. By contrast, Reo infection led to more than twenty times the number of DEG (804), again, predominantly via upregulation. Perhaps unsurprisingly, gene expression ontologies during Reo infection were almost all linked to innate immunity (Figure 5c, Figure S9).

During treatment of A549-AT cells, Reo again caused a dramatic shift in gene expression compared to the relatively quiescent SARS-CoV-2 infected cells, and this increased with the amount of time that Reo was present. The number of DEG observed following superinfection after (245), during (800), or before (1922) SARS-CoV-2 infection illustrated growing predominance of Reo-driven gene expression patterns (Figure 5d-f, Figure S10), with the pretreatment group leading to even more changes than seen for Reo infection alone. This progression was further supported by hierarchical clustering, which grouped SARS-CoV-2, untreated controls, and the 6 hpi treatment together compared to the other treatments and Reo infected cells (Figure 5g).

### Early Reo-induced ISG include novel effectors of SARS-CoV-2 restriction

We reasoned that DEG induced early during superinfection would include effectors associated with the clearance of SARS-CoV-2 infection, and that their expression would increase over time in accordance with the progressive dominance of Reo within cultures. Comparing the top 50 DEG amongst all treatment conditions rendered a list of 70+ genes that were predominantly induced at +6 hpi with expression increasing over time (Figure 6a, b, Figure S11, 12). Eliminating those already expressed within SARS-CoV-2 expressing cells produced a shortlist of ∼30 DEG (Figure 6c, Figure S12), of which around a third were predicted to form a close-knit series of interactions (Figure 6d) using STRING [23].

**Figure 6.**
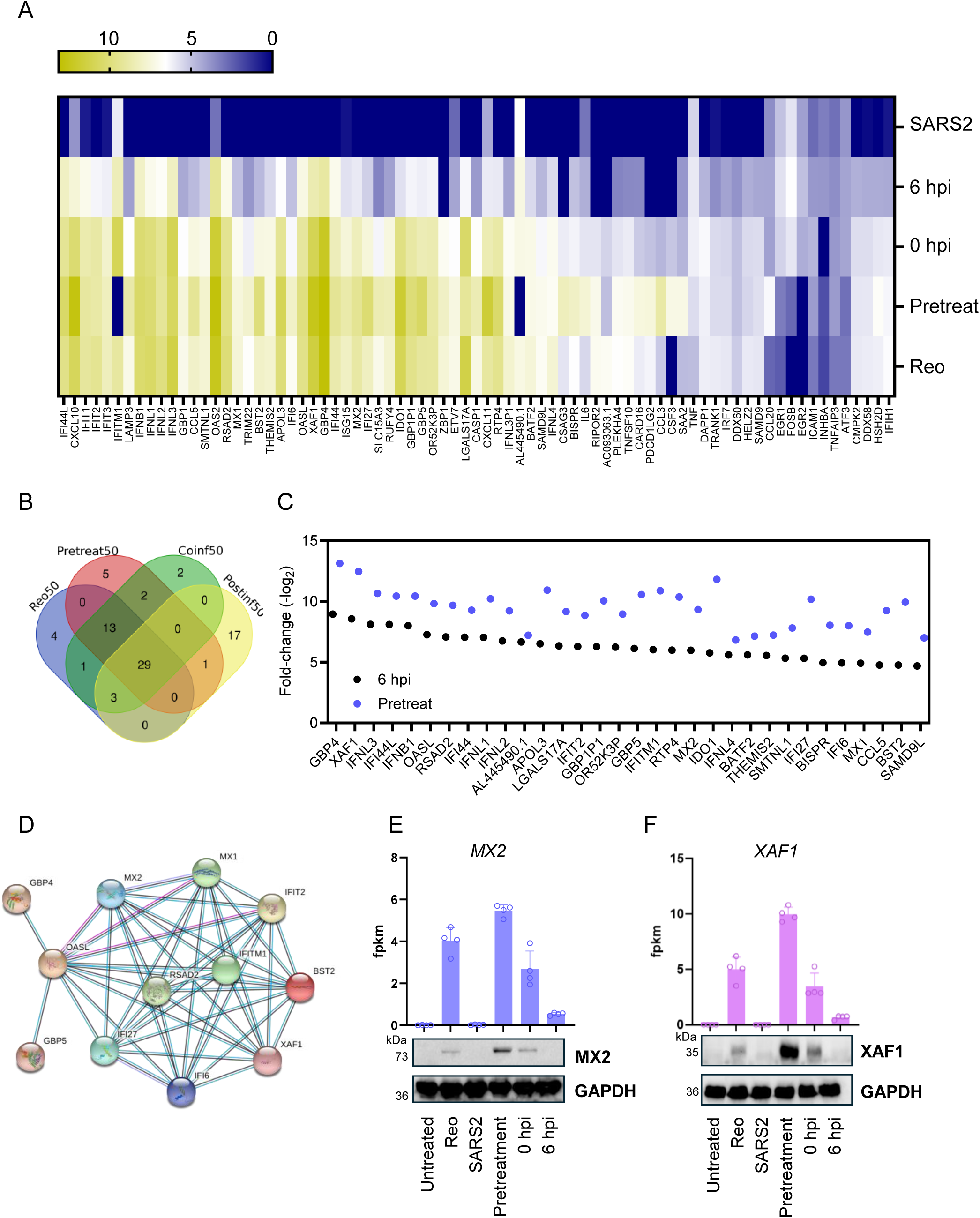
Top 50 Reo superinfection up-regulated genes are primarily ISG. **A.** Heatmap of differential gene expression for the pooled top 50 up-regulated genes during single or superinfections. **B.** Venn diagram showing distribution and overlap of DEG between data sets. **C.** Short-listed top DEG following exclusion of those already expressed within SARS-CoV-2 infected cells, showing expression at earlier (black) and later (blue) time points. **D.** High stringency String interaction analysis of shortlisted genes from C. **E-F**. Protein expression of Reo-induced MX2 and XAF1 increased in accordance with FKPM readings in RNA-Seq experiments.

We investigated whether protein expression correlated with increasing FKPM values for a subset of the gene products. Surprisingly, of eleven tested, only three exhibited protein expression patterns corresponding to mRNA levels, namely: CASP1, MX2 and XAF1 (Figure 6e, f, Figure S13). As interactions between SARS-CoV-2 and CASP1 have been extensively studied, we focused upon MX2 and XAF1 as their functional role during infection has not previously been explored.

A549-AT cells were transduced using Lentiviruses [24] either overexpressing (with RFP tag), or encoding shRNA targeting, MX2 and XAF1 (Figure 7a), along with empty vector and scrambled shRNA controls. shRNA polyclonal lines were readily selectable by puromycin resistance and prevented the induction of MX2 or XAF1 following Reo treatment (Figure 7a). By contrast, overexpressing lines took some time for RFP+ve cells to predominate, and expression levels were considerably lower than controls despite ostensibly robust expression of MX2 and XAF1 to levels exceeding that of Reo induction (Figure 7a, b). Overexpressing lines were unable to tolerate FACS enrichment for RFP expression unlike controls; we presume that overexpression reduced cell viability following stress-induced responses.

**Figure 7.**
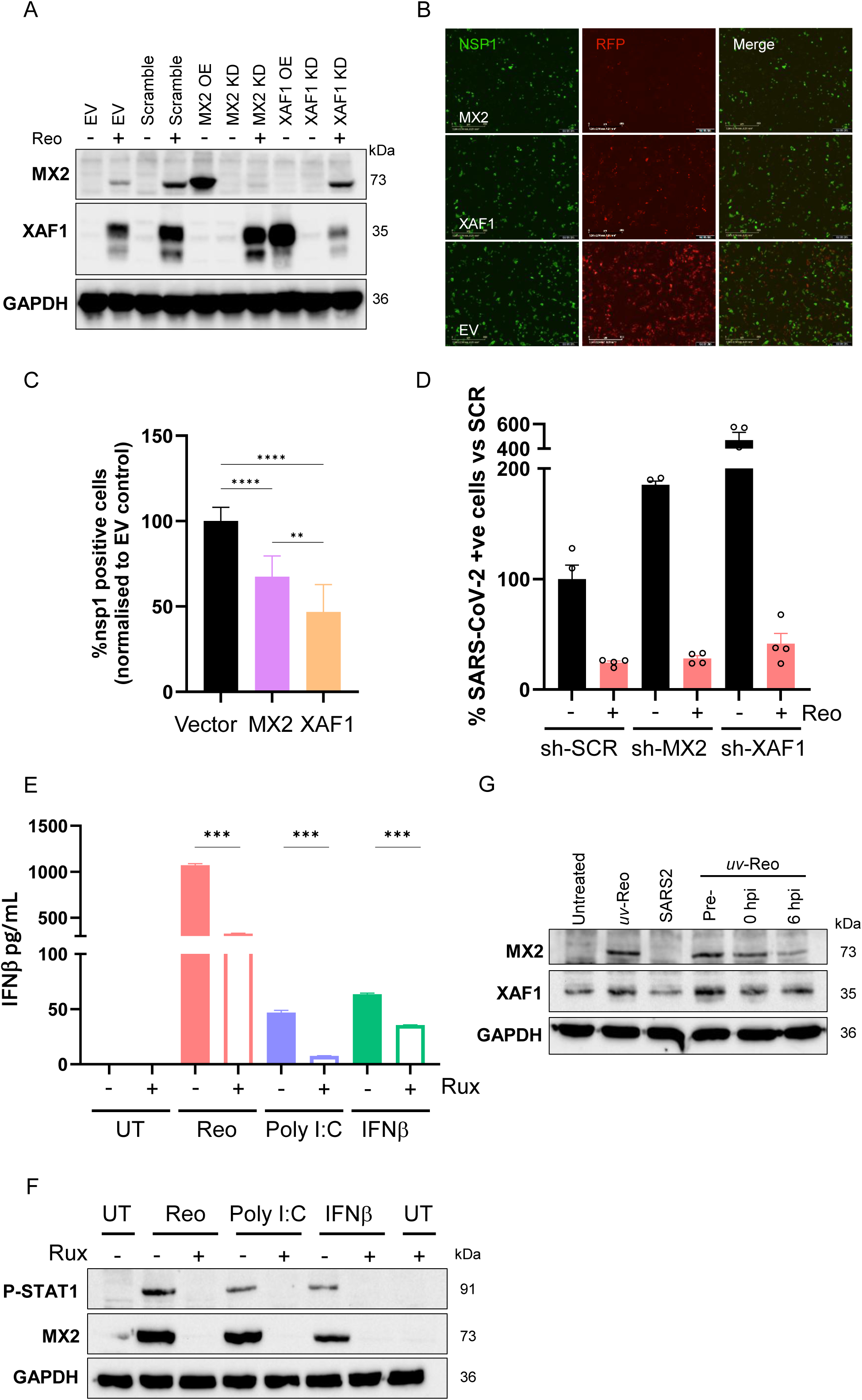
MX2 and XAF1 restrict SARS-CoV-2 replication in cell culture. **A.** Characterisation of *Lentivirus* transduced A549-AT cell polyclonal lines expressing shRNA targeting *MX2* and *XAF1*, or overexpressing them from an exogenous promotor. Western blots show expression, or lack thereof of parental and stable cell lines in the presence or absence of Reo treatment (shRNA lines only). **B.** Immunofluorescent detection of SARS-CoV-2 NSP1 (green) in polyclonal lines over-expressing MX2 or XAF1 alongside mCherry (red) from a separate cistron. **C.** Quantitation of NSP1 expression following SARS-CoV-2 infection of A549-AT cells over expressing either MX2 or XAF1. Data represent the mean ± SEM of three biological replicates and were analysed using ordinary one-way ANOVA with Tukey’s multiple comparisons test (*P < 0.05, **P < 0.01, ***P < 0.001, ****P < 0.0001). **D.** As C, but in A549-AT cells expressing MX2/XAF1 targeted-shRNAs and in the presence/absence of Reo. **E.** Comparative secretion of IFNβ measured by ELISA 24 h following stimulation of A549-AT cells with Reo, transfected Poly I:C, or IFNβ in the presence or absence of Ruxolitinib. **F.** Immunoblots of STAT-1 phosphorylation and MX2 expression within A549-AT cells corresponding to ELISA data in E. **G.** Immunoblot of A549-AT MX2 and XAF1 expression at 24 hpi following treatment of SARS-CoV-2 infected cells with *uv*-Reo.

Nevertheless, despite mosaic expression, overexpressing polyclonal populations were significantly less permissive to SARS-CoV-2 infection, compared with empty vector controls. Moreover, closer inspection revealed that the majority of NSP1+ve cells were negative for RFP, and vice versa. However, in accordance with MX2/XAF1 antiviral effects, MX2 and XAF1 shRNA knock-down lines were two- or four-fold more susceptible to SARS-CoV-2 infection, suggestive that baseline levels or low-level induction by SARS-CoV-2 for these ISGs may influence infection. Interestingly, neither cell line prevented Reo-induced suppression of SARS-CoV-2 replication, likely due to the intensity, breadth, and diversity of Reo driven ISG rewiring. Thus, MX2 and XAF1 exhibit properties consistent with SARS-CoV-2 restriction factors, in spite of issues relating to their efficient overexpression; notably, this conceivably may have prevented their effects being prominent during previous studies involving ISG expression screens.

## Discussion

The broad antiviral effects of IFN make it an excellent candidate for virus-agnostic strategies towards pandemic preparedness, or treatment of viral diseases for which no tailored therapies exist. Host IFN responses dictate the course of both acute and persistent virus infections, and numerous examples illustrate how the ongoing evolution of viruses, as well as their ability to cross species barriers, are largely determined by the dynamic between innate immunity and virus-encoded antagonists [5]. The COVID-19 pandemic has provided a real-time account of how host genetics pertinent to these interactions dictate susceptibility to severe disease, especially in the absence of prior adaptive immunity [6–8]. Viable innate-targeted immunotherapy could have made tremendous impact during the early stages of the outbreak and could continue to protect vulnerable patients, including those unable to respond to vaccines in particular. Importantly, not only does IFN target virus replication directly during the early phases of the innate response, but it links this to the establishment of effective adaptive immune responses.

Unsurprisingly, multiple clinical trials have attempted to treat both SARS-CoV-2 [15–17] and other virus infections via administering systemic recombinant IFN. These include the use of types I, II, and III IFN, although type I predominates, with regimens including both systemic delivery and inhalation of stabilised cytokines. However, despite encouraging early-stage findings, SARS-CoV-2 trials have failed to reach their primary endpoints in terms of efficacy [15, 16]. Furthermore, treatment of chronic hepatitis B (+/- D) and hepatitis C virus (HBV, HCV) infections formerly involved many months of IFN therapy, with limited success, and incurring significant toxicity that precluded many patients in need. Resolving the paradox of the preclinical effects of IFN compared to failings *in vivo* is therefore a priority.

IFNα was the first cancer immunotherapy approved by the FDA in 1986, yet despite continued use targeting both haematological (hairy cell and chronic myeloid leukaemias) and solid (melanoma, renal carcinoma) tumours, IFNs are now perceived to be obsolete [25, 26]. However, it is increasingly recognised that stimulation of endogenous IFN responses, rather than continuous use of high-dose cytokines, can lead to improved tumour outcomes; use of pattern recognition receptor (PRR) agonists and OV successfully turns immunologically “cold” tumours hot. Given that many of the immunosuppressive features within tumours resemble those produced during viral infection (e.g. IL-10, TGFβ, etc.), we investigated the utility of Reo as an innate immune adjuvant versus SARS-CoV-2 in culture to obtain proof-of-principle that Reo superinfection could rewire the innate suppression established within CoV-infected cells. Reo was chosen based upon our previous work targeting underlying oncogenic virus infections [20], as well as its inherent tissue tropism and availability as a clinical grade reagent, “Pelareorep”. Notably, this has “fast-track” FDA status for both breast and pancreatic cancer and carries an extremely favourable patient safety record.

The dsRNA Reo genome with its 5’ diphosphate is a potent RIG-I agonist [27]. Combined with the severe attenuation of type 3 Dearing strain innate antagonism, comprises a powerful innate immune stimulus. Notably, the specific laboratory strain of Reo used as Reolysin/Pelareorep (T3D^PL^, “Patrick Lee” laboratory strain) induces such potent responses far more so than viruses differing by just a handful of amino acids in key viral antagonist proteins (T3D^TD/KC^, “Terence Dermody/Kevin Coombs” labs), and this is associated with the enhanced “oncolytic” phenotype in cancer cells [28–31]. By contrast, SARS-CoV-2 infection of A549-AT cells failed to induce measurable IFN responses and RNA-Seq confirmed a minimal effect upon gene expression at 24 hpi. Notably, A549 cells express relatively low basal levels of PRR’s including RIG-I and MDA-5 (Figure 1c); Calu-3 cell basal expression is much higher (Figure S3a), but these cells are less tractable in terms of SARS-CoV-2 infection. Both PRRs sense SARS-CoV-2 RNA [32] and their functions and ensuing signalling are antagonised by multiple viral factors [33]. However, Reo superinfection led to robust A549-AT cell IFN responses, even when SARS-CoV-2 had a six-hour head-start to establish a refractory state. Ultimately, this led to a near total rewiring of gene expression towards a highly inflammatory phenotype within superinfected cultures, coincident with diminished SARS-CoV-2 infection and minimal cytotoxicity. Interestingly, stimulation with *uv*-Reo led to similar responses albeit with different kinetics, and this was Ruxolitinib-sensitive supporting a major role for paracrine IFN signalling following recognition of the Reo PAMP. Moreover, the number of coinfected cells within treated cultures was low. Accordingly, (*uv*)-RCM containing type I/III IFN as well as other cytokines effectively diminished SARS-CoV-2 replication in Vero-T cells, used due to their lack of endogenous IFN production.

The ability of Reo to override the lack of IFN response to SARS-CoV-2 during A549-AT superinfection was confirmed by RNA-Seq. This revealed a progressive switch from a proliferative/metabolic phenotype to a pronounced hyperinflammatory state resembling, but also surpassing, that evoked by Reo infection alone. Myriad ISG and IFNs were upregulated, including many already documented to restrict SARS-CoV-2 replication. However, focusing upon ISG induced at early times that continued to increase at the protein level led us to identify MX2 and XAF1 as potential new effector molecules targeting SARS-CoV-2.

XAF1 expression has been correlated with IFN responses to several RNA viruses, where it mediates IFNβ-TRAIL induced apoptosis, stabilises IRF-1 to promote ISG transcription, and also translocates to the nucleus where it regulates chromatin accessibility following TBK1 mediated Ser-252 phosphorylation at MAVS adaptor complexes [34–36]. It also acts as a tumour suppressor, driving increased p53 activation [37, 38]. However, whilst it has been shown to be increased in expression during SARS-CoV-2 infection, it has not been specifically shown to restrict viral replication [39].

Similarly, MX2 is present in multiple COVID-19 sequencing studies both from patients and preclinical systems yet has not been explicitly shown to act as a SARS-CoV-2 restriction factor [40]. MX2 is best known for its restriction of HIV-1 [41], which it mediates via both direct binding to viral capsid proteins within cytosolic condensates as well as sequestration of nucleoporins involved in their nuclear ingress [42–44]. However, MX2 has also been shown to restrict herpesviruses via similar mechanisms [44], as well as HCV via sequestration of cyclophilin A (CypA), an essential virus cofactor [45, 46]. Interestingly, MX2 and XAF1 are known to interact within cancer cells, mediating IFN induced growth arrest in melanoma [47]; in this scenario, expression of these proteins was tightly correlated, and MX2 regulated XAF1 mediated arrest via its effects upon p53. Conceivably, a similar mechanism may be at play within virus infected cells.

Overexpression of XAF1 and MX2 rendered A549-AT cells refractory to SARS-CoV-2 infection, with significant reductions in the number of infected cells detected at 24 hpi despite populations being highly heterogeneous in terms of RFP, and so presumably also MX2 and/or XAF1 expression. Conversely, shRNA mediated knock-down rendered cells more permissive, consistent with both proteins acting as SARS-CoV-2 restriction factors, either at basal levels or following low-level/transient induction following infection. However, Reo superinfection remained effective in knock-down cells, which is unsurprising given the number of reported SARS-CoV-2 effector ISGs expressed as a result. Nevertheless, the association of these factors with SARS-CoV-2 outcomes as well as new routes to understanding the fundamental biology involved warrant further investigation.

The question remains whether the observed IFN-independent induction of MX2 (or XAF1) contributes to a significant proportion of the Reo-driven anti-SARS-CoV-2 activity observed herein. MX2 is known to localise within both the peri-portal nuclear membrane as well as within cytosolic condensates. SARS-CoV-2 life cycle stages are also reported to involve these organelles, including encapsidation of the genomic RNA within condensates, and possible nucleocytoplasmic shuttling of the viral nucleoprotein (N), based upon high sequence homology with SARS-CoV-1 N. MX2 has also been shown to sequester cyclophilin A (CypA) in addition to viral capsids and nucleoporins, providing another possible mechanism by which SARS-CoV-2 might be restricted [44–46].

Whilst the findings of this study are limited by the use of cultured cell lines and their relevance to the human scenario, this is counterbalanced by cell lines allowing reliable and reproducible early-stage assessment of preclinical therapeutic efficacy. They are also highly tractable and, whilst A549-AT cells harbour lower unstimulated levels of e.g. RIGI/MDA5 expression compared with other pulmonary cell lines such as BEAS-2B, findings from ISG studies have reliably translated to the human scenario [5]. However, it will be critical to assess superinfection using primary cells and preclinical models going forwards. Future studies, including primary human epithelial models, could benefit from including further RNA-Seq analysis following RCM, IFN and *uv*-Reo treatments, as well as superinfections in the presence of IFN blocking antibodies and/or JAK inhibitors.

In summary, we demonstrate that “oncolytic” Reo superinfection may provide an alternative means of stimulating endogenous innate responses to combat pathogenic viruses, a strategy that may provide advantages over exogenous recombinant cytokines for repression of SARS-CoV-2. In addition to near complete rewiring of gene expression and a unique coinfection antiviral state, albeit *in vitro*, Reo has favourable properties that lend themselves to clinical translation and repurposing, including amplification via replication, inherent pulmonary and GI tropism, and an exemplary safety record in some of the least well patients. Its adjuvant properties are capable of undoing IFN repressive scenarios within SARS-CoV-2 infected cultures, leading to a potently antiviral state. Mechanistic differences between exogenous cytokines and Reo-induced endogenous responses will be valuable to explore in preclinical models from both a clinical and a molecular virology perspective.

## Methods

### Cell culture

Human lung-derived A549 alveolar adenocarcinoma cells overexpressing angiotensin converting enzyme 2 (ACE2) and serine protease transmembrane serine protease 2 (TMPRSS2; A549-AT), and African green monkey kidney (VeroE6) cells overexpressing TMPRSS2 (Vero-T) were obtained from the NIBSC Centre for AIDS reagents (CFAR). Calu-3 bronchial adenocarcinoma cells were purchase from Caltag Medsystems Ltd. All cells were cultured at 37 °C in a humidified atmosphere at 5 % CO_2_. A549-AT and Vero-T were cultured in Dulbecco’s modified Eagle medium (DMEM, Gibco) containing high glucose, supplemented with 10 % heat-inactivated foetal bovine serum (FBS, Gibco) and 1 % non-essential amino acids (Sigma-Aldrich). Calu-3 cells were cultured in Eagle’s Minimum Essential Medium (EMEM, Sigma-Aldrich) with the same supplements. Cells were routinely tested for Mycoplasma contamination every three months.

### Virus stocks

Clinical grade Type 3 Dearing (T3D) Strain human *Orthoreovirus* (Reo) from the Patrick Lee laboratory (T3D^PL^) was provided at specific titre by Oncolytics Biotech Inc, Calgary, CA. To generate “*uv*-Reo”, 200 µL of Reo were placed in a well of a 24 well plate and irradiated for 2 min at 254 nm *uv* light from an 8-watt bulb in a “Stratalinker” UV Crosslinker 1800 (Stratagene) as described previously [20].

SARS-CoV-2 Eng/2 (hCoV-19/England/02/2020) was obtained from Public Health England, while variants Omicron BA.2 (hCoV-19/England/FCI-179/2022) and Delta B.1.617.2 (MS066352H) were a kind gift from Rupert Beale (The Francis Crick Institute). SARS-CoV-2 stocks were prepared in-house by infecting 1.5 million Vero-T cells at a multiplicity of infection (MOI) of 0.01 PFU/cell. After 120 h of incubation, media was collected, clarified by centrifugation (500 x *g*, 5 min), aliquoted, and stored at −80 °C. Titre was determined by tissue culture 50% infectious dose (TCID_50_) assays and focus forming unit (FFU) assays.

### Plasmids

Lentiviral vector plasmids expressing MX2 and XAF1 via the human cytomegalovirus minimal immediate early promoter (CMV MIEP) and controls have been previously described [24]. Vectors expressing shRNA targeting the same factors were obtained from Sigma-Aldrich (sh-MX2:TRCN0000056713; sh-XAF1:TRCN0000134449). The psPAX2 HIV-1-based packaging helper plasmid and pMD2.G vesicular stomatitis virus glycoprotein (VSV-G) expression plasmids were gifts from Didier Trono via Addgene; plasmids #12260, #12259. Plasmids were amplified using *E. coli* (DH5α, New England Biolabs) grown at 30 ^0^C (pMD2.G at 37 ^0^C) on Luria Bertani agar plates and liquid culture, under selection with ampicillin (100 mg/mL). DNA was purified using the Qiagen Maxiprep kit, according to manufacturer’s instructions.

### Generation of, and transduction with, pseudotyped Lentiviral particles

200,000 HEK293T cells in a 6 well-plate (Corning) were transfected with 3.75 μg of psPAX2 packaging plasmid, 1.25 μg of pMD2.G, and 5 μg of vector DNA diluted in OPTIMEM, using pre-equilibrated PEI (PEI:DNA, 4:1 ratio). Briefly, the day after seeding, media was replaced with 2 mL OptiMEM media (Gibco, Thermo Fisher) before adding the reaction mix (500 μL). At 24 h post-transfection, media was replaced by supplemented DMEM. From this point on, at 48, 72, and, 96 h media was collected, pooled, clarified at 500 x *g*, 5 min, and either used or snap frozen prior to storage at −80 °C.

To generate A549 AT cells overexpressing or silenced for MX2 and XAF1, 2×10⁵ cells were seeded in 6-well plates and incubated overnight. The following day, 500 μL of Lentiviral stock (vectors encoding MX2, XAF1, or shRNA targeting them, see above) and polybrene (8 μg/mL; Santa Cruz Biotechnology) were added. Cells were centrifuged using a plate spinner (1000 × *g*, 1 h, room temperature) and then incubated for 24 h at 37 °C, after which the medium was replaced with supplemented DMEM. At 48 h post-transduction, the medium was replaced with puromycin-containing medium (0.5 μg/mL) and cells passaged until negative controls succumbed to the antibiotic.

### Reo Conditioned Media (RCM)

A549 or Calu3 cells were exposed to live or *uv*-Reo (M.O.I. of 1 PFU/cell) and incubated for 24 or 48 h respectively. Media was collected, filtered using Viresolve NFP (Merck), and stored in 1 mL aliquots for long-term storage at −20 °C. Anti-SARS-CoV-2 activity of RCM was assessed using Vero-T cells infected at 0.1 PFU/cell, with treatments applied at −12 h, 0 h, or 6 h post-infection (hpi), followed by fixation with 4 % (w/v) PFA for 30 min prior to assessment of SARS-CoV-2 by focus forming assay at 24-72 hpi.

### Direct Reo and SARS-CoV-2 coinfection

A549 AT cells were infected with SARS-CoV-2 (M.O.I. of 0.5 PFU/cell), with live or *uv*-Reo added as pretreatment (−12 h), at 0 h, or 6 h post-infection at MOIs of 1, 2, or 5 PFU/cell. Cells were incubated for 24–72 h before fixation and measuring of SARS-CoV-2 infected cells by focus forming assay as above.

### Immunofluorescence (IF) and focus formation assays

Cells were fixed with prewarmed room temperature 4 % w/v PFA/PBS for 20 min, then washed in PBS twice prior to permeabilisation using 0.1 % Triton X-100/PBS (Alfa Aesar) for 10 min at room temperature. Following three further PBS washes, cells were incubated for 30 min with 5 % BSA (w/v) in PBS. Primary antibodies, anti-σNS (Reo) mouse monoclonal antibody (DSHB Hybridoma Product 2A9; 1:50), anti-nsp1 (SARS-CoV-2) sheep monoclonal antibody (NIBSC CFAR; 1:500), anti-N protein (SARS-CoV-2) sheep monoclonal antibody (NIBSC CFAR; 1:500), were applied overnight at 4 °C. After further washing, cells were incubated with secondary antibodies diluted 1:200-500 in 5 % BSA/PBS as above (Invitrogen) and/or Hoechst (Molecular Probes; 1:10,000), specifically: goat anti-mouse Alexa Fluor 488 nm, donkey anti-goat Alexa Fluor 488 nm, chicken anti-mouse 594 nm, donkey anti-sheep Alexa Fluor 555, chicken anti-mouse Alexa Fluor 647, donkey anti-rabbit Alexa Fluor 488, for 1 h in the dark at room temperature. Imaging checks were performed using an EVOS FL microscope (Thermo Fisher Scientific), with quantification of infected cells on an IncuCyte S3 system (Sartorius) as described previously. High resolution co-stained images were taken using Zeiss LSM 980 106 confocal microscope.

### Western blotting

Cells were harvested and lysed using Mammalian Protein Extraction Reagent (M-PER, Thermo Fisher Scientific) for BSL2 samples (Reo), or RIPA (Radioimmunoprecipitation assay: 150 NaCl, 50 mM Tris pH 7.4, 0.5% Sodium Deoxycholate, 0.1% SDS, 1 % NP40, 1mM EDTA, 1% Triton X-100 and dH_2_O) buffer for BSL3 samples. Both were supplemented with protease and phosphatase inhibitor tablets (Thermo Fisher Scientific). Protein concentration was determined using a Pierce BCA Protein Assay Kit against a BSA standard curve (Thermo Fisher Scientific), and 30 μg loaded per well. Samples were adjusted using 4x Laemmli buffer (BioRad) and ∼1.43 mM β-mercaptoethanol (Sigma), then heated at 95 °C for 5 min. Proteins were separated on 10 % tris-glycine SDS-PAGE gels, then wet transferred to PVDF (polyvinylidene difluoride) Immobilon FL transfer membranes (Merck Millipore, IPFL00010) in transfer buffer (25 mM Tris base, 192 mM glycine, 0.1 % (w/v) SDS, and 20 % (v/v) methanol) for 1 h 30 min – 2 h at 120 V. Membranes were washed three times (for 5 min) with Tris-buffered saline (TBS) with 0.1 % (v/v) Tween 20 detergent (TBS-T) and blocked for 15 min with 5 % (w/v) Bovine Serum Albumin (BSA) (Fisher Bioreagents) dissolved in TBS-T. Primary antibody (Cell Signalling Technology (1:1000): rabbit anti-RIG-I (3743), rabbit anti-MDA5 (5321), rabbit anti-Ser536 phospho-NFkB p65 (3033), mouse anti-NFkB p65 (6956), rabbit anti-Ser396 phospho-IRF3 (29047), rabbit anti-IRF3 (4302), rabbit anti-Tyr701 phospho-STAT1 (9167), rabbit anti-STAT1 (9172). Santa Cruz Biotechnology (1:100): Mouse anti-MX2 (sc-271527), mouse anti-XAF1 (sc-398012)) previously diluted in 5 % BSA/TBS-T was added and incubated overnight at 4 °C. The following morning, the membrane was TBS-T washed three times for 5 min, and then an appropriate horseradish peroxidase labelled secondary antibody was added diluted (1:5000) in 5 % BSA/TBS-T (Sigma Aldrich: Goat anti-mouse A4416, Goat anti-rabbit A6154) and incubated for 1 h. After three further rounds of washing, Super Signal West Pico Plus Chemiluminescent Substrate (Thermo Fisher Scientific) was used to detect and quantify signal from antibody-labelled proteins using a ChemiDoc MP Imaging System (BioRad).

### Cytokine array

Cytokine profiling of live or *uv*-Reo A549 RCM was performed using the Human Cytokine Antibody Array C5 (RayBiotech) according to the manufacturer’s instructions, with CM and serum-containing DMEM as controls. Signals were detected using the ChemiDoc MP Imaging System, and dot intensities were quantified via densitometry in ImageJ.

### Enzyme-linked immunosorbent assay (ELISA)

Quantitative detection of secreted IFNβ and IFNα were performed using the Human IFN-beta Quantikine ELISA Kit and the Human IFN-alpha (α) All Subtype Quantikine ELISA Kit (R&D Systems). Type III IFN was measured using Human IL-28B/IFN-lambda 3 ELISA Kit (RayBiotech) and Human IL-29 ELISA Kit (Invitrogen). Optical density was measured at 450 nm on a Cytation 5 Imaging Plate Reader.

### RNA extraction and sequencing

A549 AT cells were coinfected with SARS-CoV-2 Eng/2 (0.5 PFU/cell) and Reo (5 PFU/cell), along with both single and control mock infections. Total RNA was extracted 24 h post-infection using TRIzol (Thermo Fisher Scientific) following the manufacturer’s protocol, with *n* = 4 biological replicates per condition. RNA quality was assessed by optical density at 230, 260, and 280 nm on a Nano Drop spectrophotometer, and samples were sent to Novogene for additional quality control (Agilent 5400) prior to sequencing.

Libraries were prepared using poly-T magnetic bead mRNA isolation, constructed and quality checked, and sequenced on an Illumina NovaSeq 6000 (150 bp paired-end reads). Clean reads were aligned to the human reference genome (GRCh38/hg38) with HISAT2, and expression levels were estimated as FPKM (Fragments Per Kilobase of transcript sequence per Million base pairs sequenced).

Three-dimensional principal component analysis (3D-PCA) indicated excellent separation of clustered biological repeats, confirmed by Pearson Correlation. FPKM distribution was similar, and a homologous median and interquartile range existed between samples and groups. Differential expression was assessed with DESeq2 (|log₂FC| > 1, padj < 0.05), followed by gene enrichment analysis (Novagene), and overlapping gene lists were visualised using <u>Venn</u> <u>diagrams</u>. Raw sequencing data will be available following upload to the Gene Expression Omnibus (GEO) genomics data repository.

## Acknowledgements

We are grateful to Oncolytics Biotech (Calgary, CA), and its past members, for generous provision of clinical grade reovirus stocks for research. We are also grateful to those generously providing plasmids and other reagents, as listed in the methods. This work was supported by the Secretariat of Science, Humanities, Technology and Innovation (SECIHTI, Mexico) (SGC), and MRC grant MR/T016205/1 (awarded to SG).

## Supplementary Figure Legends

**Figure S1.**
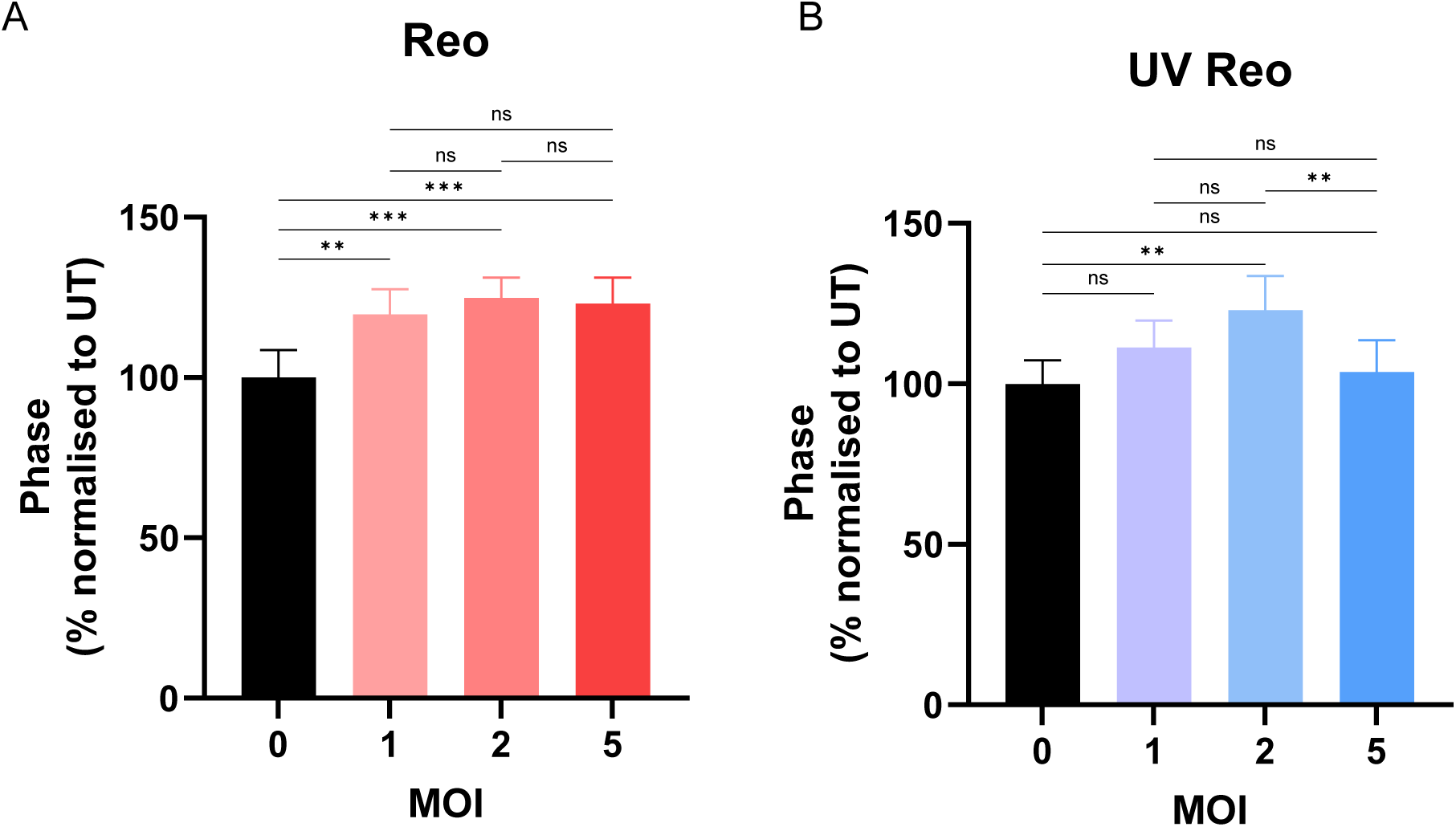
Viability of A549-AT cells treated with increasing MOI of **A.** Reo and **B.** *uv*-Reo, measured by % confluency using the IncuCyte Zoom.

**Figure S2.**
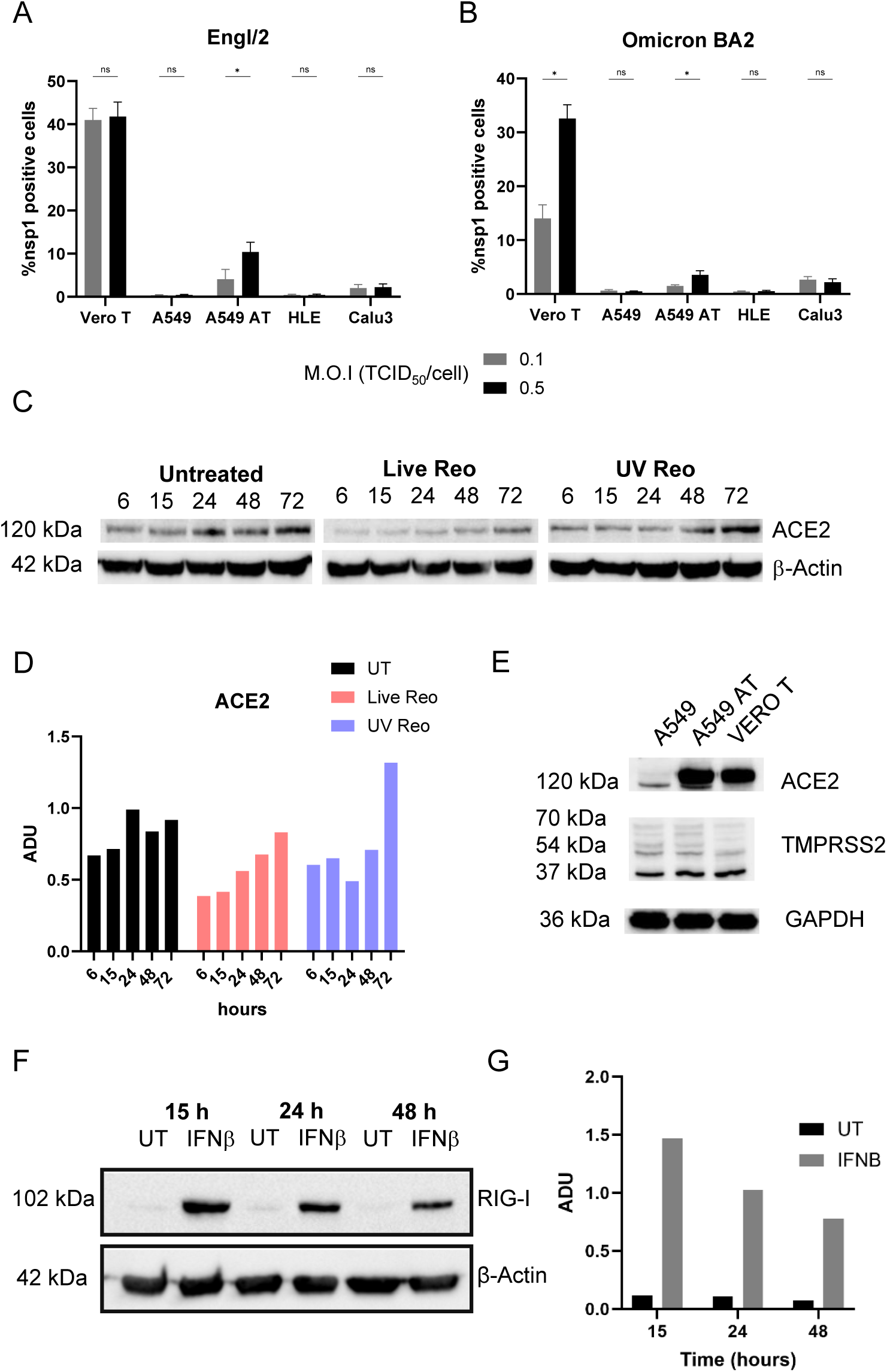
Characterisation of SARS-CoV-2 experimental infection. **A.** % NSP1 positive cells measured at 24 hpi following infection of Vero-T, A549, A549-AT, HLE, and Calu3 cells were infected with Eng/2 SARS-CoV-2 at two MOI (0.1 and 0.5 PFU/cell). Data represent the mean ± SD of three biological replicates and were analysed using multiple Mann-Whitney tests (ns = not significant, *P < 0.05). **B.** As for A but using SARS-CoV-2 Omicron BA.2. **C.** Expression of ACE2 in A549-AT cells over time comparing controls with Reo and *uv*-Reo treatment. **D**. Quantification of immunoblots shown in C. **E.** Baseline expression of ACE2 and TMPRSS2 in A549, A549-AT and Vero-T cells. **F.** Induction of ISG protein expression (RIG-I) in A549-AT treated with recombinant IFNβ. **G.** Quantification of immunoblot in F.

**Figure S3.**
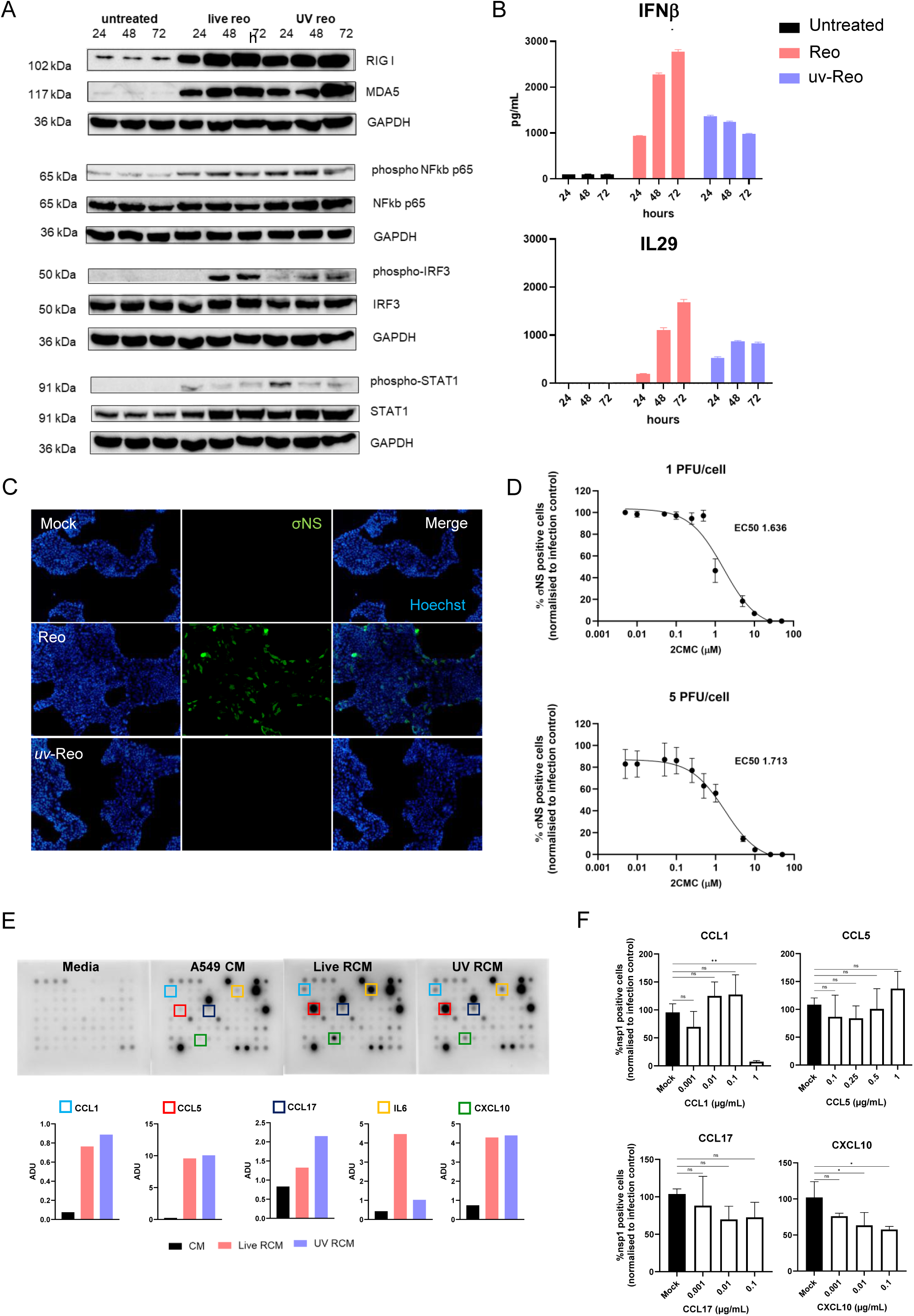
**A.** Immunoblots comparing Reo versus *<u>uv</u>*-Reo stimulation of Calu-3 cells, testing for the same pathways as per figure 1c. **B.** ELISA characterisation of (*uv*)-RCM derived from Calu-3 cells, as per figure 1d. **C.** Immunofluorescence for Reo σNS in Calu-3 cells stimulated with Reo or *uv*-Reo compared to controls. **D.** Log_2_-fold titration of 2CMC in A549-AT cells infected with Reo at a multiplicity of 1 (top) or 5(bottom) PFU/cell. Calculated EC_50_ values are shown. **E.** Raw image data for cytokine/chemokine protein arrays conducted upon (*uv*)RCM versus controls. **F.** %NSP1 positive A549-AT cells at 24 hpi with SARS-CoV-2 treated with increasing concentrations of recombinant cytokines. Data represent the mean ± SD of three experimental replicates and were analysed by using ordinary one-way ANOVA with Tukey’s multiple comparisons test (*P < 0.05, **P < 0.01, ***P < 0.001, ****P < 0.0001).

**Figure S4.**
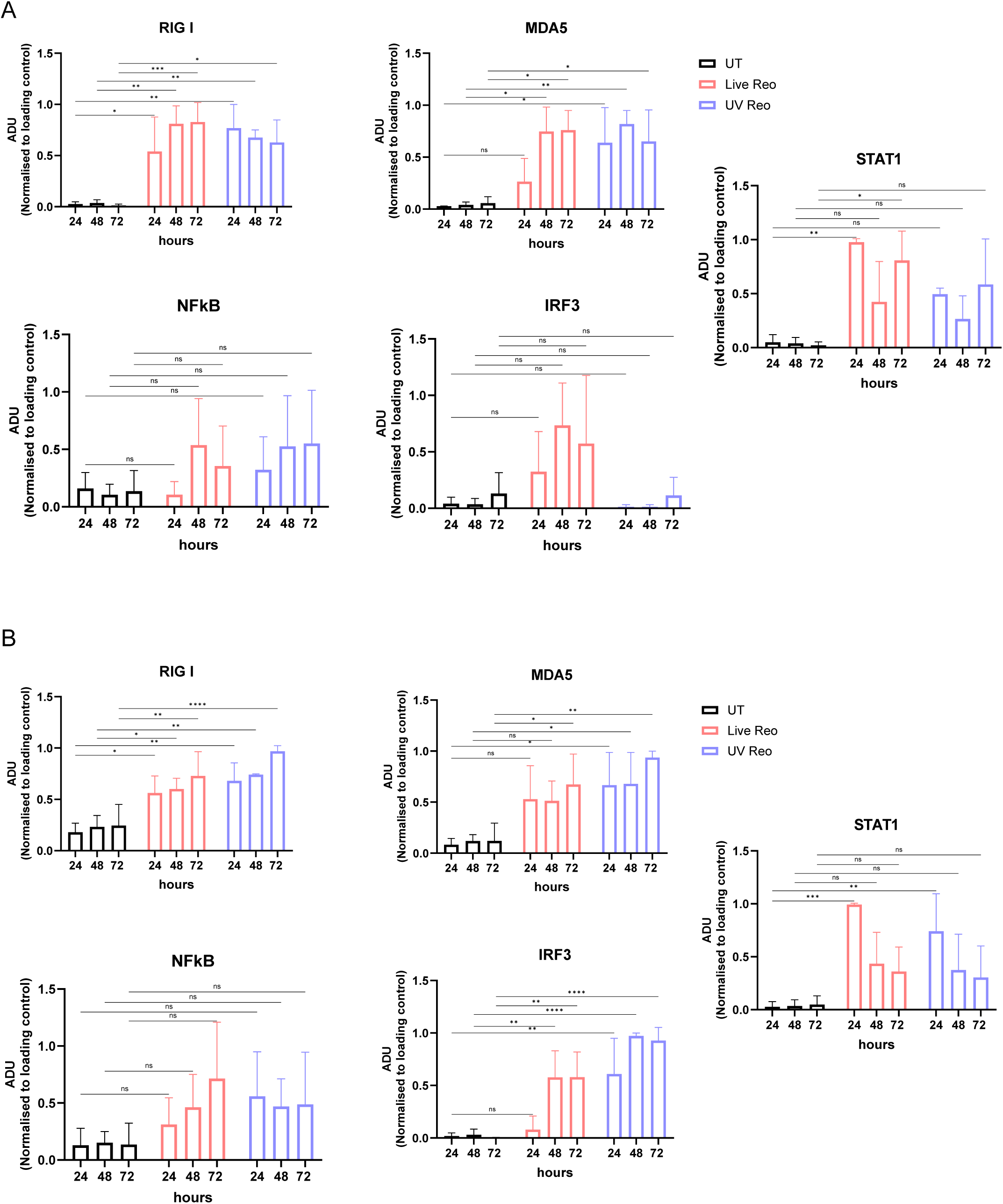
A. Quantification of immunoblots shown in figure 1c. **B.** Quantification of immunoblots shown in figure S3.

**Figure S5.**
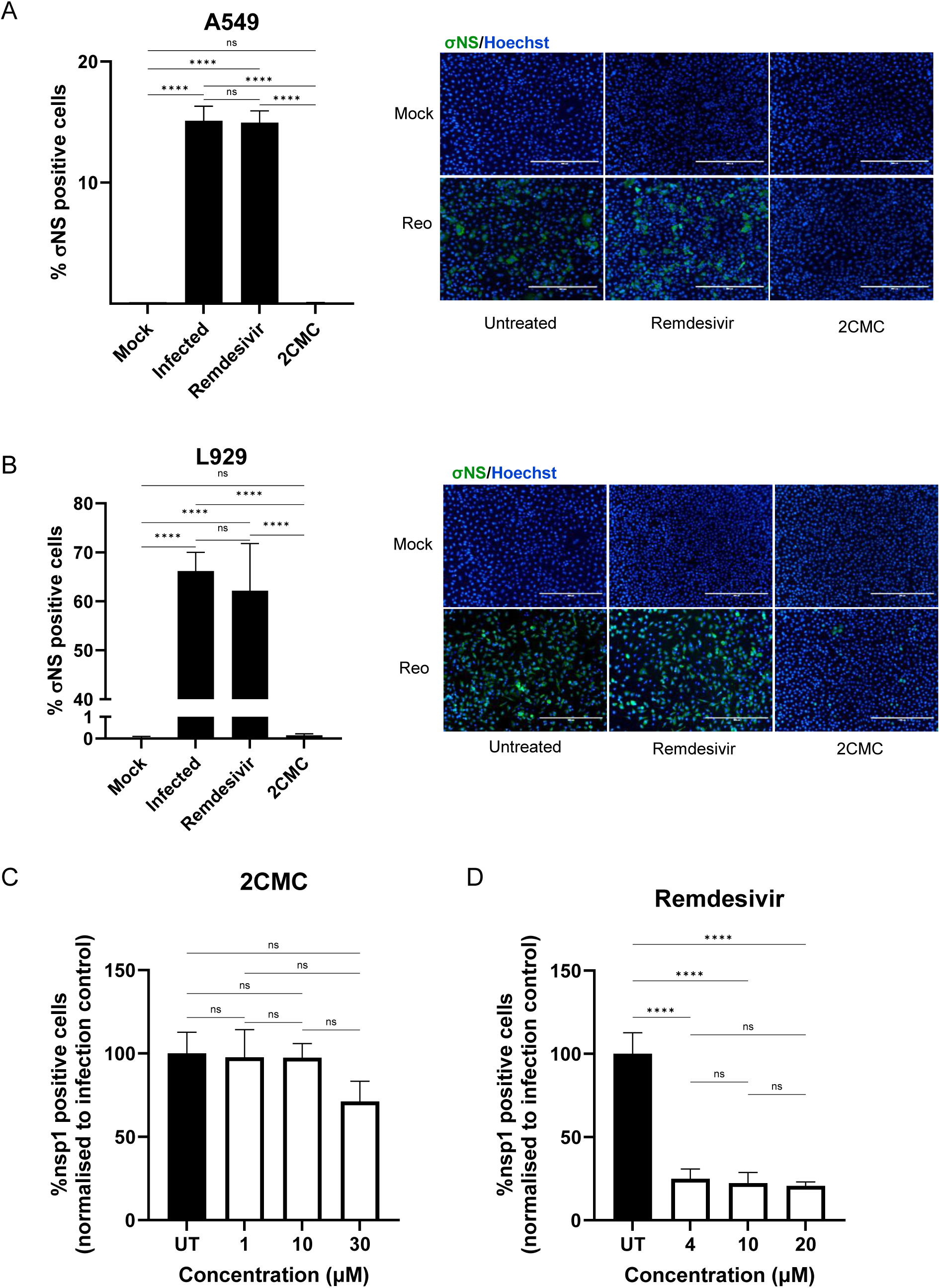
**A.** Comparative activity of 2CMC or remdesivir versus Reo replication within A549-AT cells. **B.** As A, but in highly permissive murine L929 cells. **C.** Titration of 2CMC versus SARS-CoV-2 infected A549-AT cells, measuring 24 hpi using the IncuCyte. **D.** As C, but using Remdesivir.

**Figure S6.**
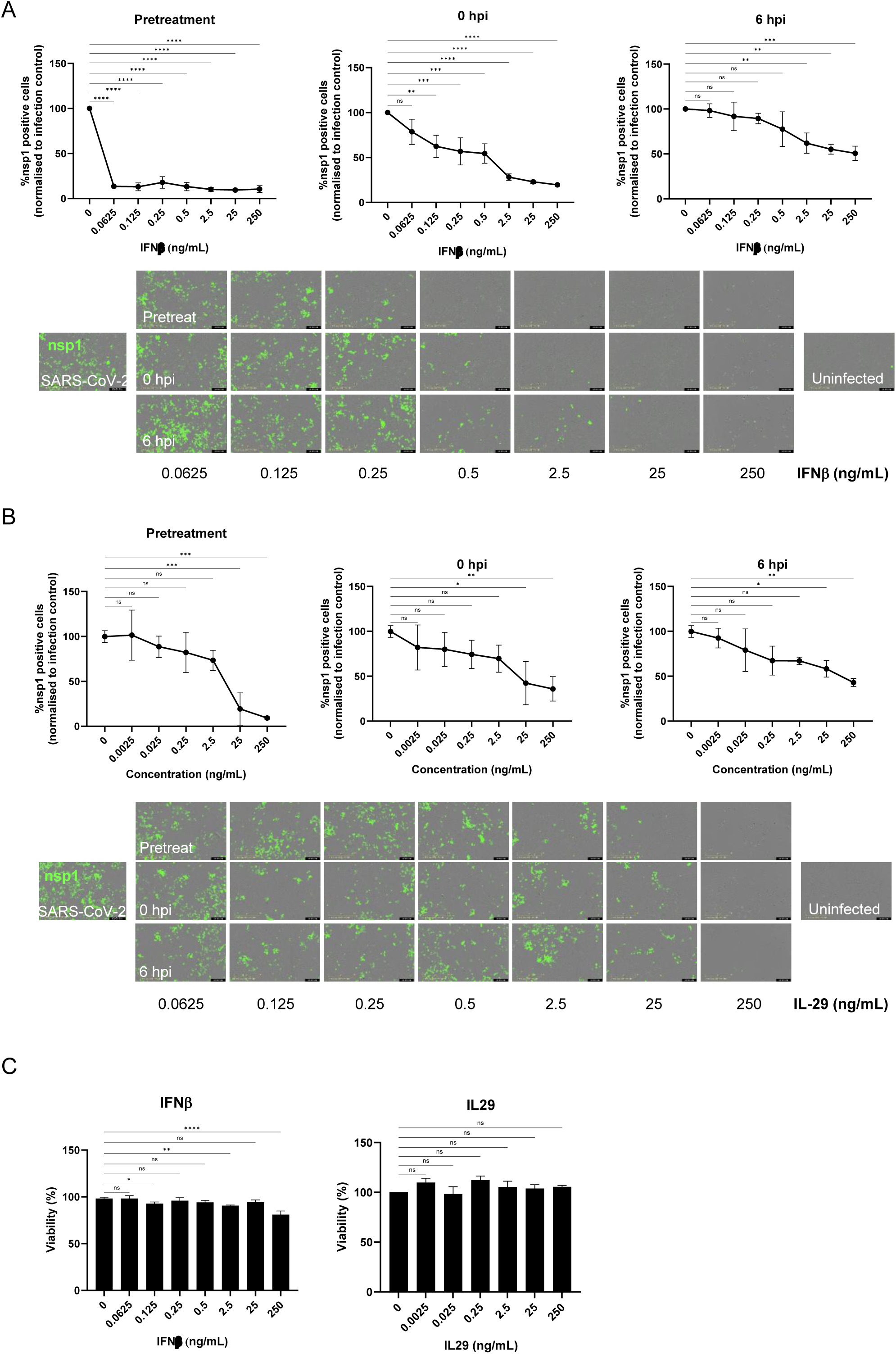
**A.** Titration of recombinant IFNβ (0.0625 - 250 ng/mL) against SARS-CoV-2 infected A549-AT cells measuring %NSP1 +ve cells at 24 hpi using the IncuCyte, cytokine added at same time points as Reo superinfection. **B.** As A, but for recombinant IL-29. **C.** Cell viability (uninfected) during cytokine titrations measured by cell confluency using the IncuCyte. Data represent the mean ± SD of three biological replicates and were analysed using ordinary one-way ANOVA with Tukey’s multiple comparisons test (*P < 0.05, **P < 0.01, ***P < 0.001, ****P < 0.0001).

**Figure S7.**
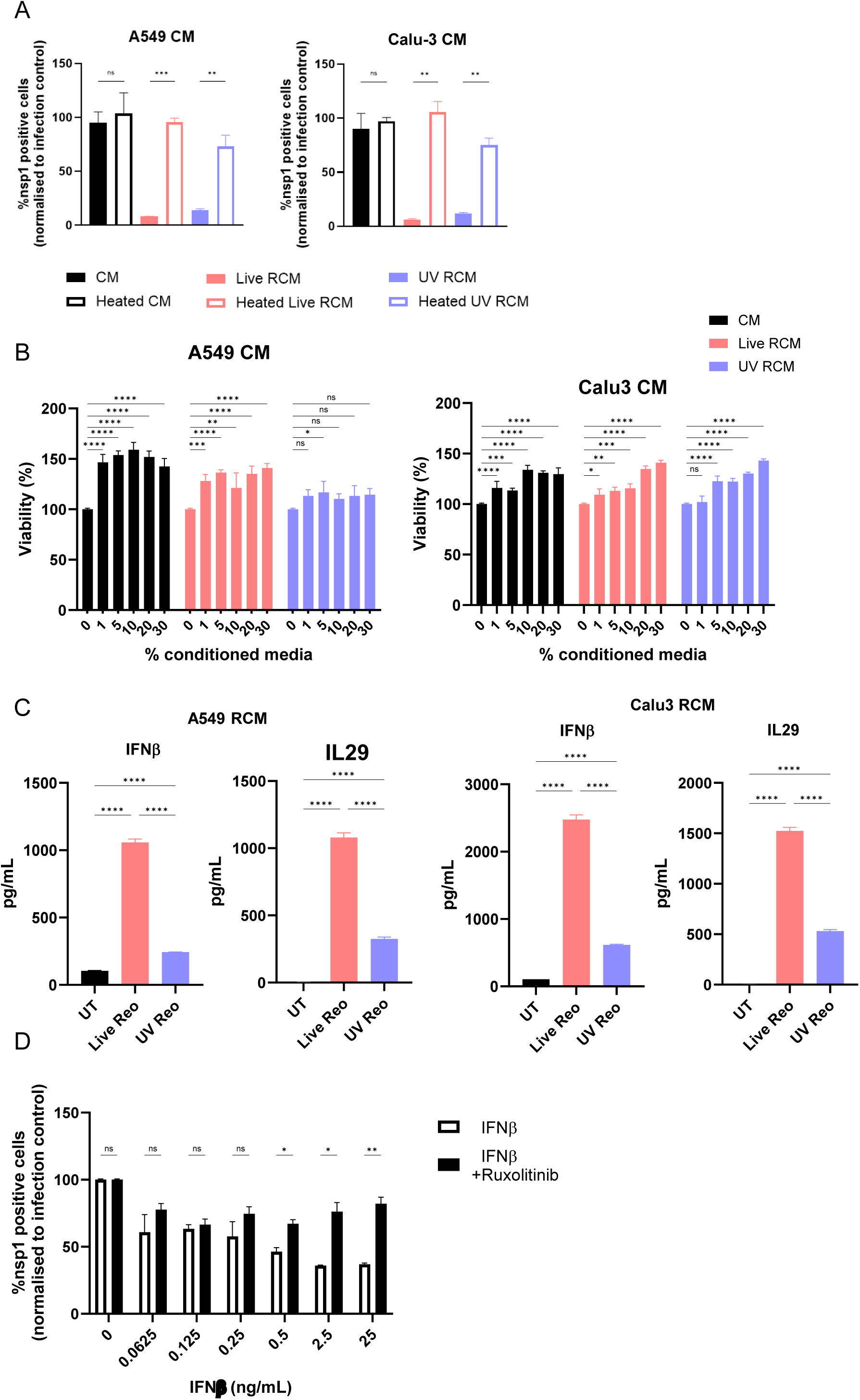
**A.** Effect of heating (90 ^0^C, 10 min) upon SARS-CoV-2 antiviral activity of conditioned media, measuring %NSP1 +ve cells at 24 hpi using the IncuCyte. Data represent the mean ± SD of three biological replicates and were analysed by ordinary multiple t-tests (*P < 0.05, **P < 0.01, ***P < 0.001, ****P < 0.0001). **B.** Vero-T cell viability 24 h following treatment with increasing concentrations of CM, RCM, or *uv*-RCM derived from A549-AT or Calu-3 cells, filtered using Viresolve NFP. Data represent the mean ± SD of three biological replicates and were analysed using ordinary two-way ANOVA with Tukey’s multiple comparisons test (*P < 0.05, **P < 0.01, ***P < 0.001, ****P < 0.0001). **C.** Comparison of IFNb and IL-29 secretion within (*uv*)-RCM derived from A549-AT (left) or Calu-3. Analysis as in B. **D.** Effect of Ruxolitinib upon SARS-CoV-2 antiviral effects in Vero-T cells treated with increasing concentrations of recombinant IFNβ, measuring %NSP1 +ve cells at 24 hpi using the IncuCyte. Data represent the mean ± SD of three biological replicates and were analysed using ordinary multiple t tests (*P < 0.05, **P < 0.01, ***P < 0.001, ****P < 0.0001).

**Figure S8.**
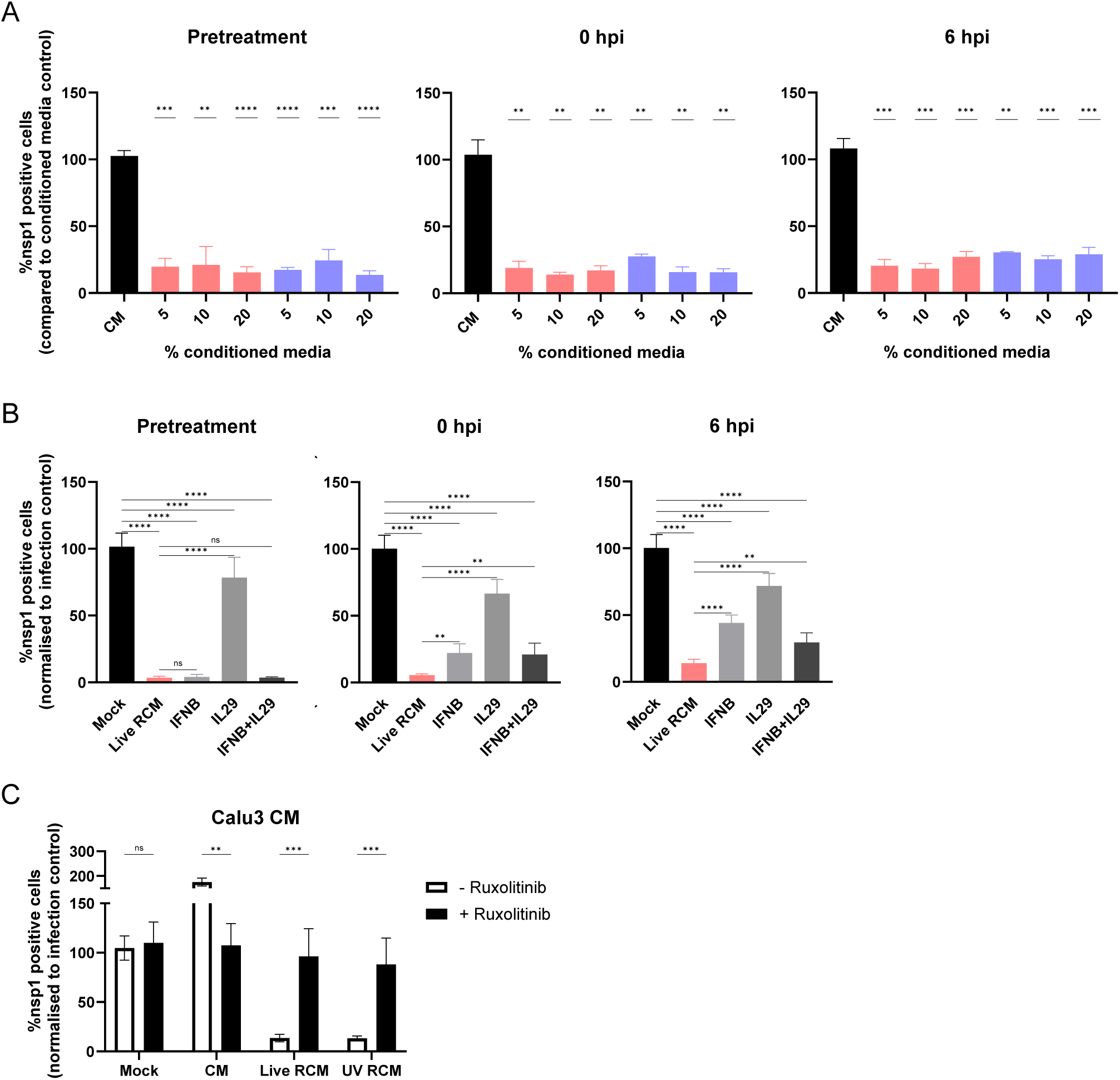
Parallel experiments conducted using Calu-3 derived (*uv*)-RCM as shown in figure 4c, 4d, and 4f, corresponding to **A, B**, and **C**.

**Figure S9.**
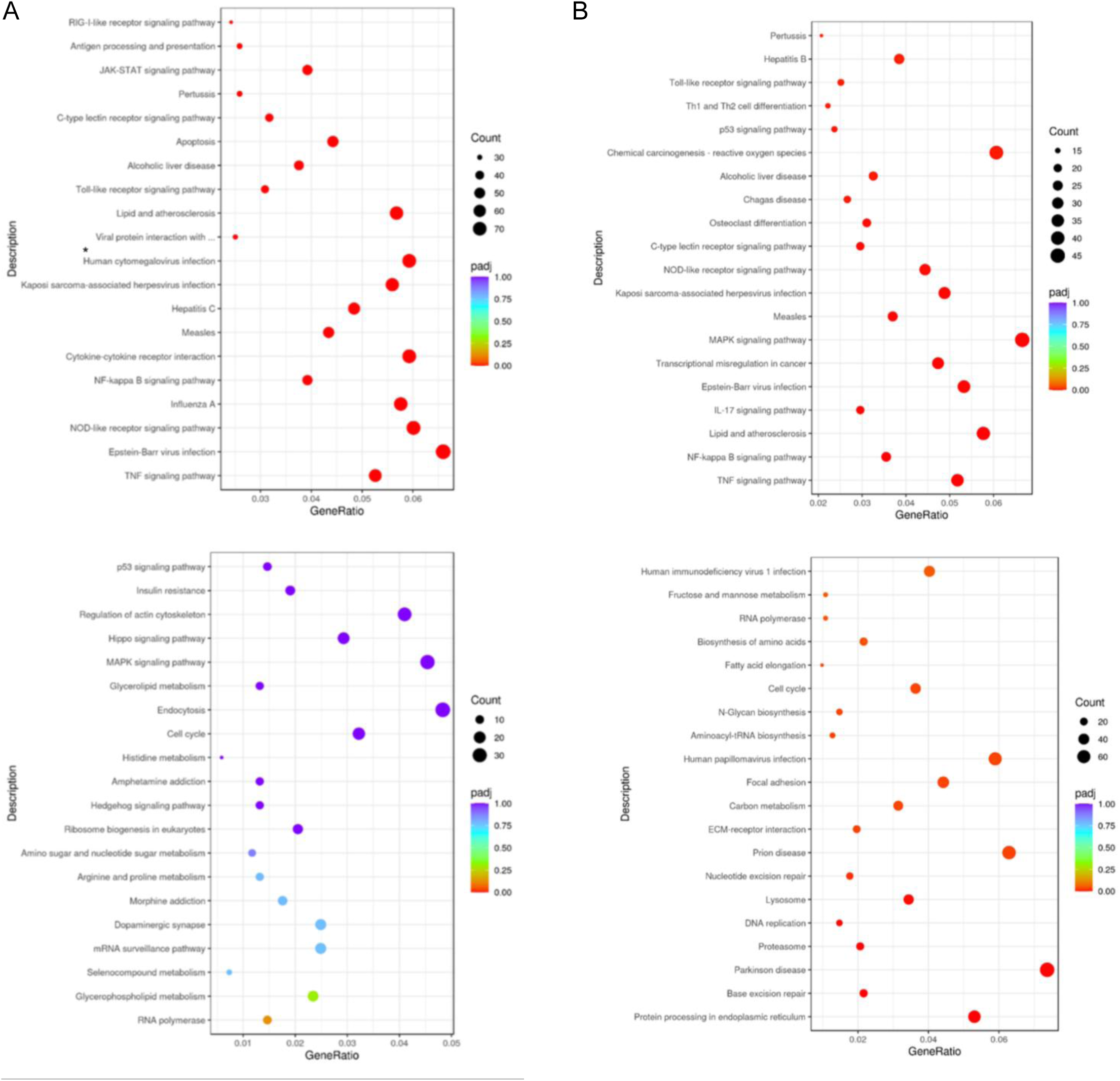
**A.** Gene enrichment analysis of differentially expressed genes (DEG) using KEGG for Reo infected A549-AT cells compared to uninfected controls. Top panel shows up-regulated, bottom down-regulated DEG. **B.** As A for SARS-CoV-2 infection.

**Figure S10.**
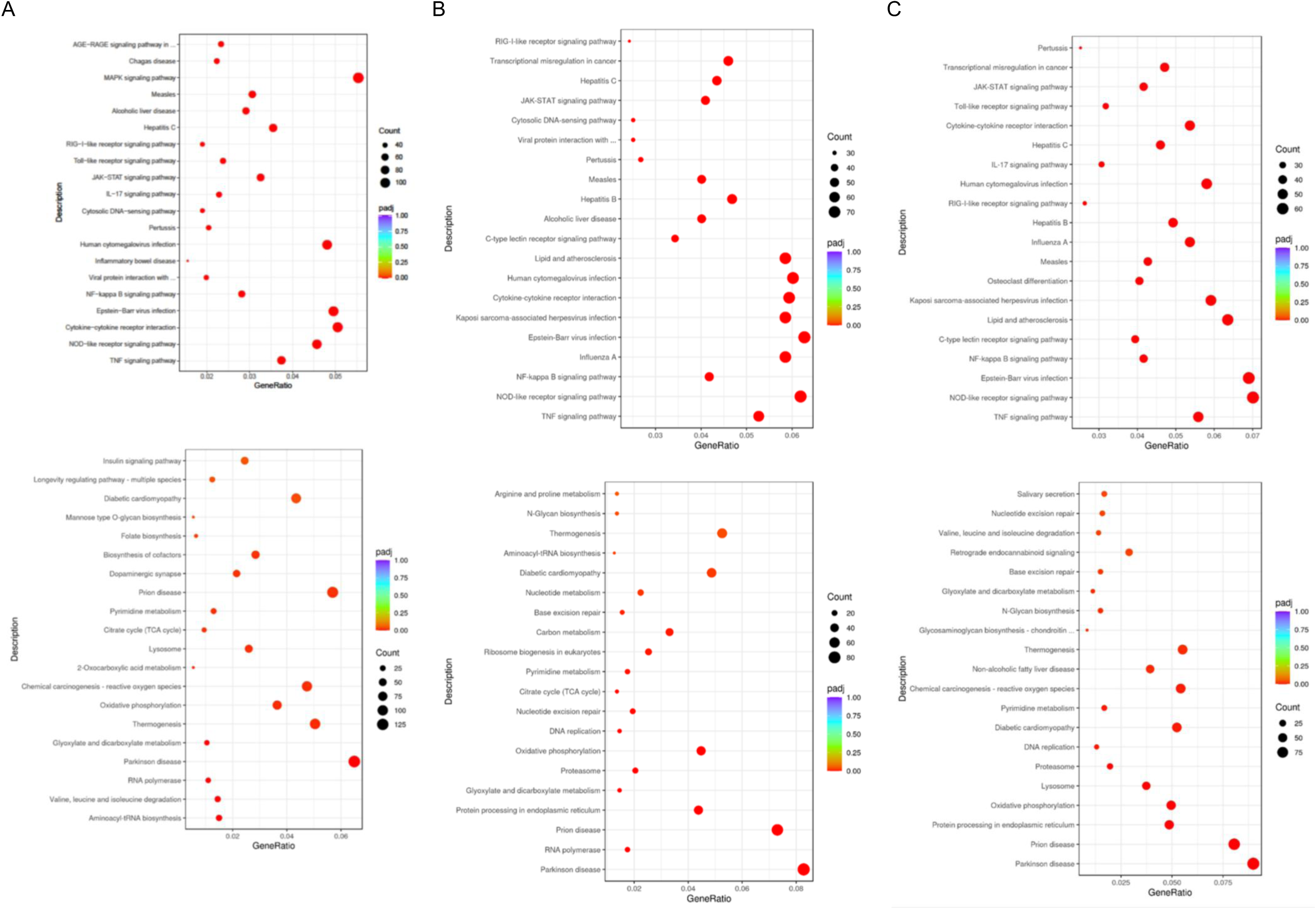
As for figure S9, but for superinfected cells 24 hpi with Reo pretreatment, 0 hpi, and 6 hpi shown in **A, B,** and **C,** respectively.

**Figure S11.**
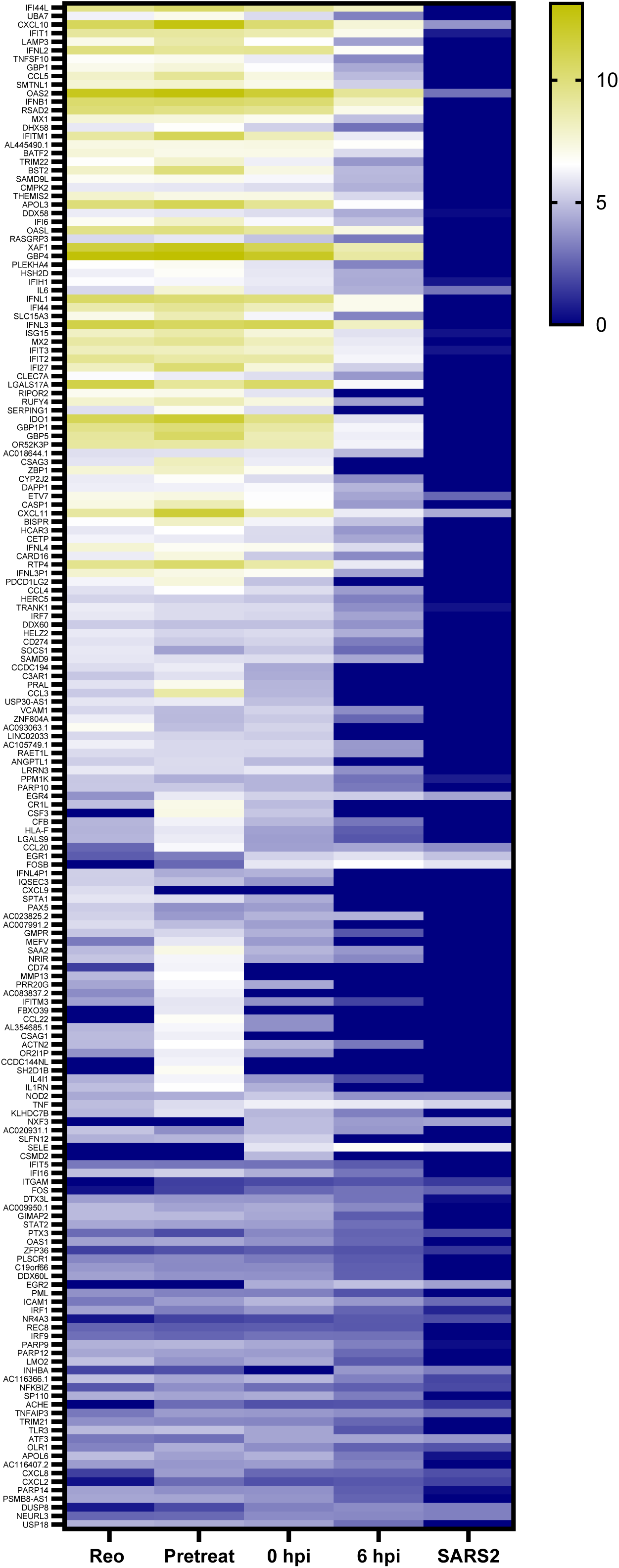
Heatmap of extended pooled “top 100 genes” up-regulated during superinfection conditions.

**Figure S12.**
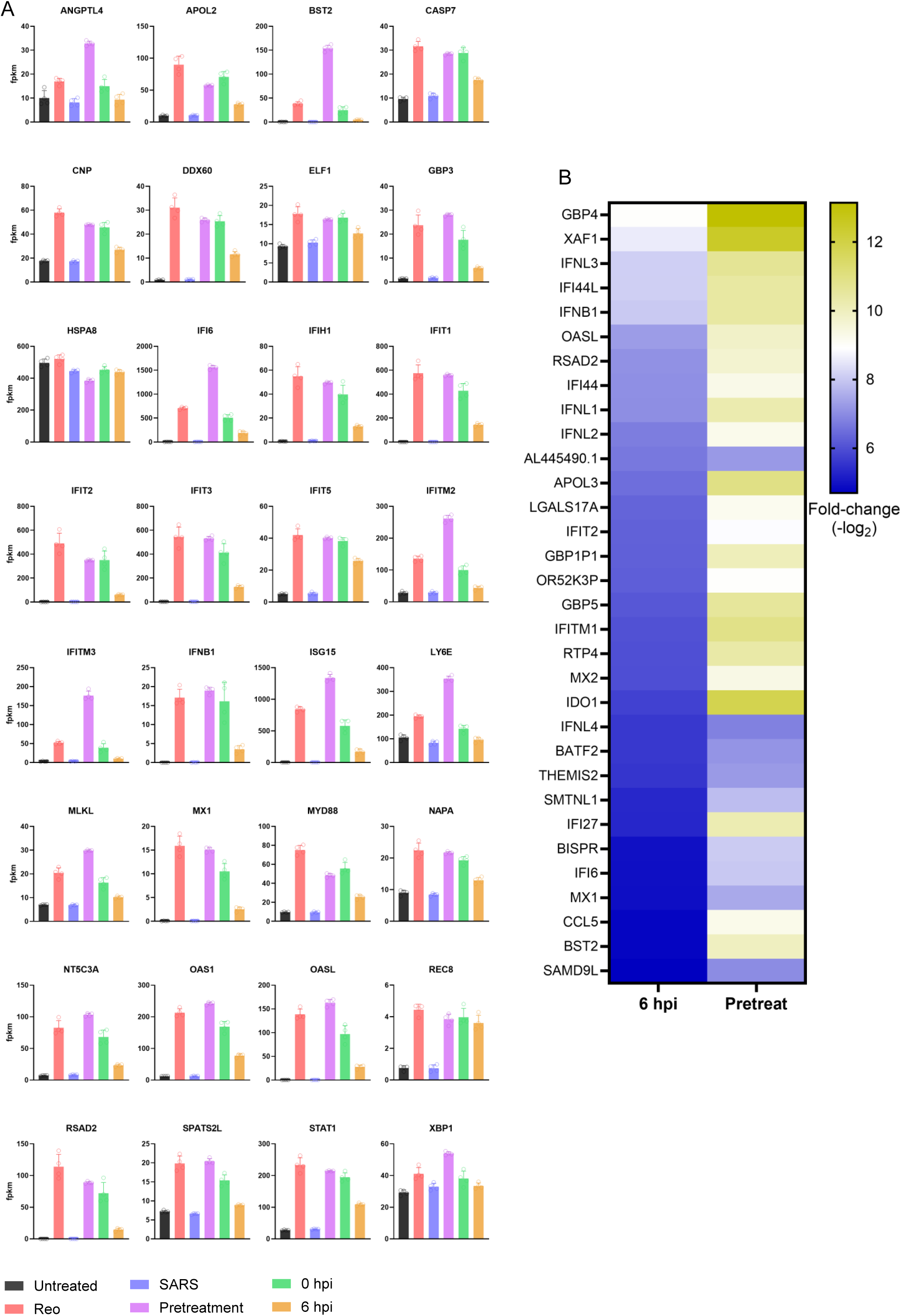
**A.** FKPM values for selected DEG expressed at early times (6 hpi). **B.** Heatmap showing refined ISG set after elimination of those expressed within SARS-CoV-2 infected cells, based upon same data as shown in figure 6c.

**Figure S13.**
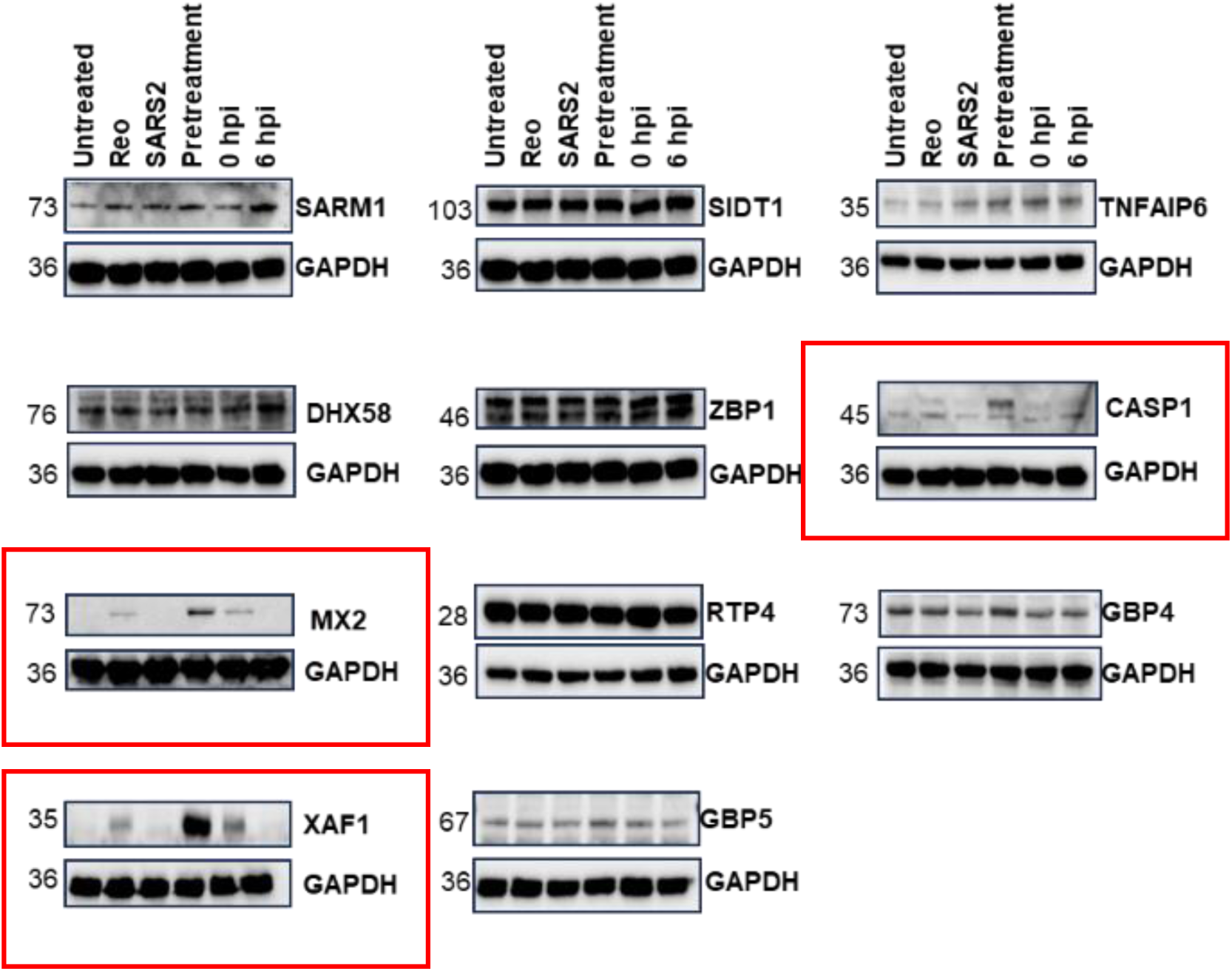
Immunoblots for selected “early ISGs” including MX2 and XAF1.

